# Functional Specialization of Parallel Distributed Networks Revealed by Analysis of Trial-to-Trial Variation in Processing Demands

**DOI:** 10.1101/2022.04.20.488923

**Authors:** Lauren M. DiNicola, Oluwatobi I. Ariyo, Randy L. Buckner

## Abstract

Multiple large-scale networks populate human association cortex. Here we explored the functional properties of these networks by exploiting trial-to-trial variation in component processing demands. In two behavioral studies (N=136 and N=238), participants quantified strategies used to solve individual task trials that spanned remembering, imagining future scenarios, and various control trials. These trials were also all scanned in an independent sample of functional MRI participants (N=10), each with sufficient data to precisely define within-individual networks. Stable latent factors varied across trials and correlated with trial-level functional responses selectively across networks. One network linked to parahippocampal cortex, labeled Default Network A (DN-A), tracked scene construction, including for control trials that possessed minimal episodic memory demands. To the degree a trial encouraged participants to construct a mental scene with vivid imagery and awareness about spatial locations of objects or places, the response in DN-A increased. The juxtaposed Default Network B (DN-B) showed no such response but varied in relation to social processing demands. Another adjacent network, labeled Frontoparietal Network B (FPN-B), robustly correlated with trial difficulty. These results support that DN-A and DN-B are specialized networks differentially supporting information processing within spatial and social domains. Both networks are dissociable from a closely juxtaposed domain-general control network that tracks cognitive effort.

## Introduction

Diverse higher-order functions, including autobiographical memory, spatial navigation and social inference, have been attributed to a large monolithic network known as the default network (DN; Buckner & Carroll 2007, Buckner et al. 2008, Spreng et al. 2009; see also Gusnard & Raichle, 2001, Hassabis & Maguire 2007, Schacter et al. 2007, Binder et al. 2009). This network extends into rostral temporal and prefrontal association cortex leading to its description as the apex higher-order association network (Margulies et al. 2016, Buckner & DiNicola 2019). Considerable attention has been given in recent years to understand the processing contributions of the DN to human cognition (e.g., Murphy et al. 2018, Sormaz et al. 2018, Beaty et al. 2020, Lee et al. 2021, Wen et al. 2021, Yeshurun et al. 2021, Mancuso et al. 2022).

One challenge for understanding processes supported by the DN is that most prior studies rely on group-averaged data, which necessarily blurs anatomical details (see Steinmetz & Seitz 1991, Fedorenko et al. 2010, Laumann et al. 2015). Recent explorations within intensively scanned individuals reveal that the DN comprises at least two fully distinct, parallel networks (Braga & Buckner 2017, Buckner & DiNicola 2019; see also Deen & Friewald 2021). These networks, termed DN-A^1^ and DN-B, contain features of previously-hypothesized DN subsystems (e.g., Andrews-Hanna et al. 2010, Yeo et al. 2011) but are fully distinct (Braga & Buckner 2017), raising questions about functional differentiation. Both networks possess regions distributed across multiple association zones with side-by-side juxtapositions throughout the cortex, sometimes on opposite sides of the same sulcus (Braga et al. 2019). Given these spatial arrangements, unraveling their distinct processing contributions has been hampered by spatial averaging over individuals.

Supporting functional heterogeneity, within-individual analyses suggest that DN-A is preferentially recruited by tasks targeting episodic remembering and imagining the future, and DN-B by tasks targeting social inferences. Contrasting these task domains reveals a replicable double dissociation (DiNicola et al. 2020; see also Rosenbaum et al. 2007, Andrews-Hanna et al. 2014, Kurczek et al. 2015). Adding further evidence for domain specialization, separate regions of DN-A and DN-B within the posterior midline differentially respond to spatial versus social content (Peer et al. 2015, Silson et al., 2019, see also Woolnough et al. 2020, Deen & Friewald 2021).

However, there is a second complicating factor for understanding the processing contributions of these juxtaposed networks. The tasks that elicit activation responses in these networks are often complex, involving temporally-extended trial structures that encourage rich and varied mental constructions (e.g., Hassabis & Maguire 2009, Schacter et al. 2012). In this sense, much like the spatial blurring that has led to ambiguities in the existing literature, the mixing of multiple task components in the lengthy task trials that activate the DN also leads to ambiguities. Open questions thus remain about the nature of the underlying processes that these recently identified, parallel networks support, as well as those that differentiate DN-A and DN-B from other juxtaposed networks. Answering questions about one’s past, for example, involves multiple processes traditionally associated with episodic memory retrieval (e.g., Tulving, 1983), as well as component processes that might generalize beyond episodic memory, such as constructing a scene in a spatially-coherent context (e.g., Hassabis & Maguire 2007, Hassabis & Maguire 2009) and deployment of domain-general controlled processing resources (Kopelman 1991, Dobbins et al. 2002, Bunge et al. 2004, Vatansever et al. 2021; see also Moscovitch 1992, Badre & Wagner 2007).

Here we explored network functions using a behavioral approach to probe trial-to-trial variation in processing demands across a diverse set of previously scanned task trials targeting episodic remembering and imagining the future. The approach did not assume specific relations between component processes and individual network responses, but allowed relations to emerge to the degree that trial-to-trial variation in processing demands selectively associated with network responses. For each trial, participants were asked questions that encouraged them to remember or imagine distinct scenarios (see examples in Figures 1 and 2). The questions were designed to vary in self-relevance (Self or Non-Self) and temporal orientation (Past, Present or Future), and afforded considerable opportunity to adopt varied strategies. Prior analyses of these functional MRI data focused on predetermined contrasts between conditions that grouped many trials together (DiNicola et al. 2020). Plotting network responses across separate trials within each condition revealed large signal variations well beyond that expected by measurement error, reinforcing that there is unaccounted for trial-level variation and creating a novel experimental opportunity.

**Figure 1.**
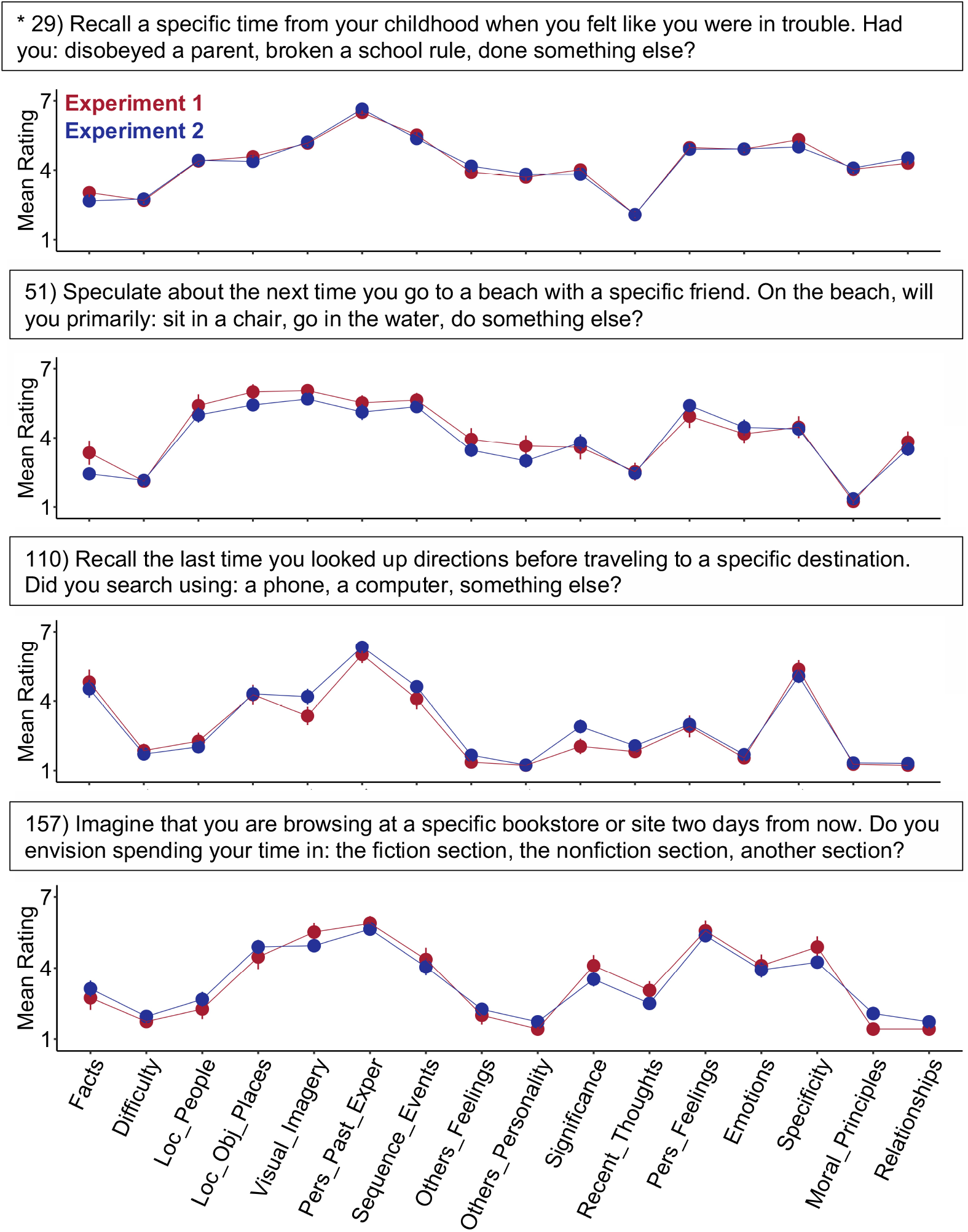
Behavioral ratings illustrate unique and reliable strategy use patterns. Mean strategy ratings from independent groups of behavioral participants show striking similarity (red: Exp. 1, blue: Exp. 2). Four example trials are displayed, chosen from the original ‘target’ conditions designed to demand episodic projection (e.g., remembering and imagining the future). Above each plot is the actual question the participants viewed; below is the measured strategy pattern. The strategies plotted on the x-axis are listed in Table 1. The trials share high ratings for strategies relevant to episodic memory, as intended, such as consideration of the personal past (Personal_Past_Exper), events (Sequence_Events) and mental scenes (Visual_Imagery, Loc_Obj_Places). High intertrial variability on other strategy dimensions highlights the exploratory opportunity (see also Figure 2). For example, Difficulty was low for some trials and higher for others. Each point shows a mean strategy rating across participants with standard error bars. Pers = Personal; Exper = Experiences. * Denotes a repeated trial with a larger sample size.

**Figure 2.**
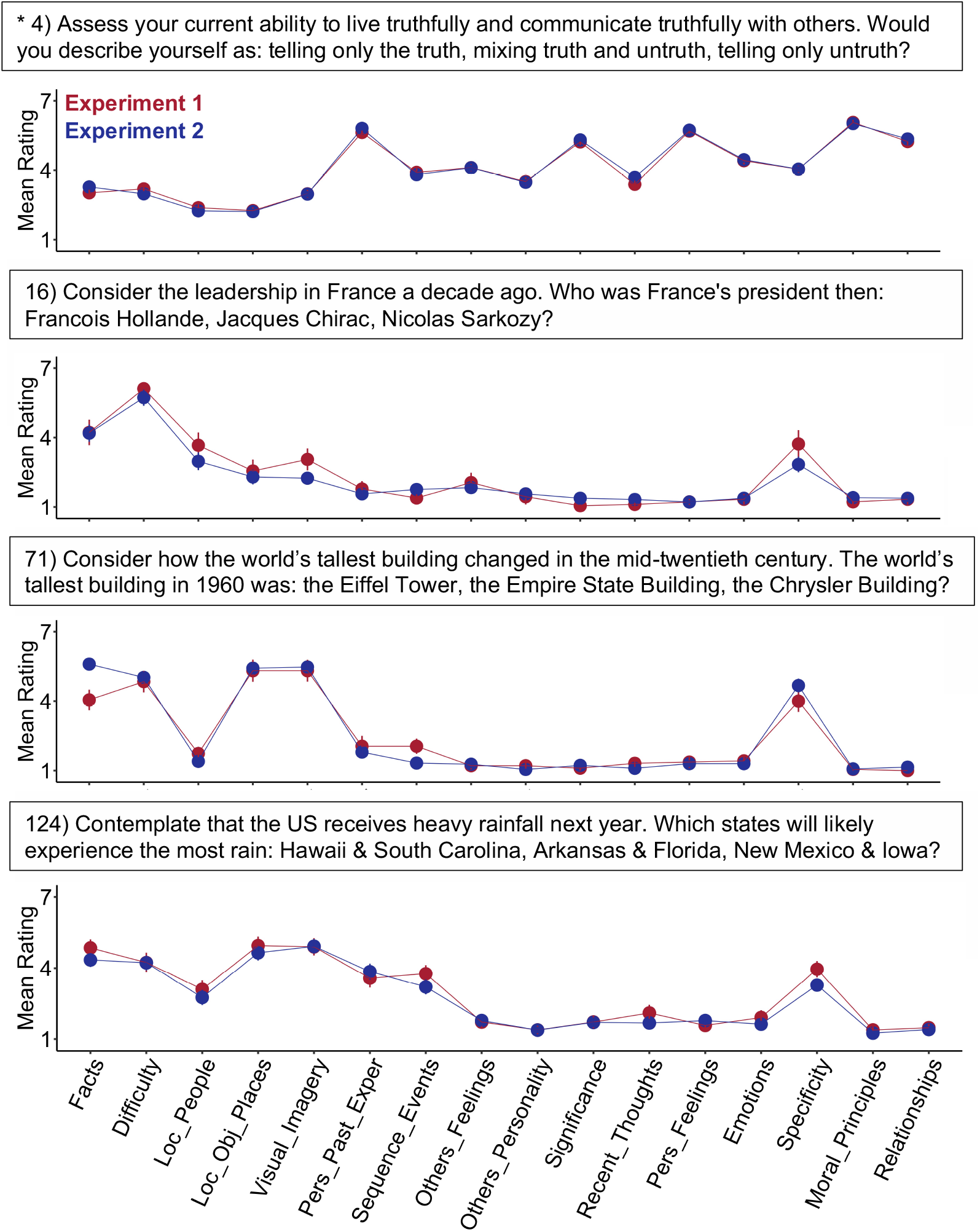
Strategy use patterns differ markedly among the control trials. Strategy patterns are shown for four trials taken from the control conditions. For each trial, mean strategy ratings from independent groups again reveal notable overlap (red: Exp. 1, blue: Exp. 2). The four trials were selected from the original ‘control’ conditions, designed to minimize demands on episodic projection. Most of these trials show lower reliance on the personal past (Pers_Past_Exper) and greater use of facts than the target trials in Figure 1. The control trials also reveal marked variability. Multiple trials involve strategies related to mental scenes (Visual_Imagery, Loc_Obj_Places), for example, or to considering unfolding events (Sequence_Events). Each point shows a mean strategy rating across participants with standard error bars. Pers = Personal; Exper = Experiences. * Denotes a repeated trial with a larger sample size.

Specifically, we behaviorally assessed *how* people responded to each individual trial question, to quantify trial-level properties for comparison to network activity (as indirectly measured by functional MRI). Evidence from prior studies support that assessing *how* people respond to complex stimuli can provide insight into processing demands (e.g., related to memory encoding – Kirchhoff & Buckner, 2006; emotion discrimination – Skerry & Saxe 2015; and differentiating physical from emotional pain - Bruneau et al. 2013). Most directly relevant to the present study, Andrews-Hanna et al. (2010) collected strategy ratings from both scanned participants and an independent behavioral group, and found that composites of strategy ratings tracked activity in network regions of interest. Strategy ratings can, therefore, tap into stable properties of individual task trials, providing an experimental approach to explore component processes. We adopted such an approach here to functionally dissociate multiple juxtaposed networks that were precisely measured within individual participants, and also to provide insight into each network’s functional contributions to task processing.

## Methods

### Overview

Data for analyses came from previously collected neuroimaging participants (DiNicola et al. 2020) paired with newly collected behavioral data that measured the idiosyncratic processing demands of each trial. The task neuroimaging data included 180 trials where participants answered unique questions by selecting one of three possible choices. In the present work, independent online behavioral participants rated the strategies they used, to assess how each of the 180 questions were answered (Table 1). The strategy ratings were remarkably stable between independent behavioral samples and could be clustered into intercorrelated composites. Functional properties of the networks were examined by asking whether activity levels in distinct brain networks preferentially tracked strategy composite scores.

**Table 1.** The Response Strategies Scale (RSS) included 16 strategy probes.

| # | Abbreviation | Strategy Probe | Mean Rating | SD Rating | Inter-Experiment Reliability (*r*) |
| --- | --- | --- | --- | --- | --- |
| 1 | Significance | **To what extent did this question** ... ask about a matter that is significant to you? | 3.18 | 1.59 | 0.92 |
| 2 | Pers_Feelings | ... lead you to think about your own preferences, feelings or emotions? | 3.63 | 1.59 | 0.93 |
| 3 | Emotions | ... evoke any emotion (e.g., happiness, sadness, excitement, etc.)? | 3.02 | 1.62 | 0.91 |
| 4 | Pers_Past_Exper | ... rely on personal past experiences? | 4.15 | 1.71 | 0.93 |
| 5 | Sequence_Events | ... lead you to imagine a sequence of events unfolding in your mind? | 3.61 | 1.86 | 0.91 |
| 6 | Loc_Obj_Places | ... lead you to envision the location of objects or places mentioned in it? | 3.70 | 1.98 | 0.89 |
| 7 | Loc_People | ... lead you to envision the physical locations of other people? | 2.83 | 1.82 | 0.86 |
| 8 | Others_Feelings | ... lead you to speculate about the preferences, emotions, or thoughts of other people? | 2.90 | 1.75 | 0.92 |
| 9 | Moral_Principles | ... make you consider general moral principles? | 1.89 | 1.25 | 0.91 |
| 10 | Relationships | ... require you to think about the nature of your relationships to other people? | 2.16 | 1.30 | 0.96 |
| 11 | Others_Personality | ... lead you to consider the personality traits or appearances of other people? | 2.28 | 1.58 | 0.88 |
| 12 | Visual_Imagery | ... evoke visual imagery in your mind? | 4.30 | 1.89 | 0.81 |
| 13 | Recent_Thoughts | ... evoke thoughts that have been on your mind recently? | 2.49 | 1.53 | 0.91 |
| 14 | Difficulty | **When answering this question** ... how hard did you have to think? (1 – thoughts were spontaneous or easy, 7 – thoughts were deliberate or difficult) | 3.02 | 1.65 | 0.93 |
| 15 | Facts | ... to what extent did you rely on facts, as opposed to subjective experiences? (1 – relied only on subjective experiences, 7 – relied only on facts) | 3.85 | 1.95 | 0.86 |
| 16 | Specificity | ... how specific were your thoughts? (1 – thoughts were broad and general, 7 – thoughts were detailed and specific) | 4.21 | 1.91 | 0.66 |

Notes: Two phrases ('To what extent did this question...' and 'When answering this question...', bolded in the table) were each followed by multiple probes. All probes included a 7-point scale (for probes 1-13: 1 – None, 7 – A lot; for 14-16: scales shown in the table). Strategy probes 1-8, 14 and 15 were adapted from Andrews-Hanna et al. (2010; see their Table S3). For each probe, rating means and standard deviations (SD) are shown, calculated across raters in Exp. 1 and Exp. 2. Inter-experiment reliability shows the Pearson's correlation between mean ratings in Exp. 1 and Exp. 2, across all trials, for each probe.

### Experiment 1: Initial Exploration of Strategy Ratings

#### Participants

175 paid participants ages 18-27 were recruited from Amazon Mechanical Turk using Cloud Research (Litman et al. 2017) to answer trial questions and complete surveys about their strategy use while answering the questions. Participants were English-speakers within the United States who had high ratings for completion of prior studies (90%+ approval rating for at least 100 prior tasks). Each participant provided informed consent through an online protocol approved by the Institutional Review Board (IRB) of Harvard University. Question and strategy probe forms were created and administered using Harvard’s Qualtrics platform.

#### Question and Strategy Probe Format

Online participants answered questions from the episodic projection task of DiNicola et al. (2020). After each individual question, the participants scored the strategies they used across 16 probes designed to tap into a variety of possible processing components (Table 1). Given the burden of rating many strategy probes, each participant answered only a subset of the original trial questions (34-38 total trials per participant). The trials were rotated across participants so that 25 participants rated strategies used for each unique trial. As in the original task, each trial asked a question about either a real-world experience or general knowledge, and featured three answer choices (see DiNicola et al. 2020). A few trials required minor wording changes for generalization to online participants (e.g., changing “a trip out of Boston” to “a trip out of your town”). Finally, as control procedures, a subset of trials was repeated, including 5 shown to all participants (each from a different condition) to test for potential cohort effects, as well as two attention checks (i.e., one that probed whether participants were reading questions and another targeting task focus).

Trial timing was as follows: participants first saw the question and answer options, and were asked to respond within 10 sec, mirroring the neuroimaging protocol. After 10 sec, participants received a reminder to select a response. After answering the question, the participants were then presented with the Response Strategies Scale (RSS; Table 1) and asked to rate their use of 16 strategies on a scale from 1 to 7. The RSS expanded upon the scale used by Andrews-Hanna and colleagues (2010) to incorporate previously-assessed strategies (e.g., reliance on memory, personal significance, effort), as well as new strategies, informed by work probing mind-wandering and memory components (e.g., consideration of people’s attributes, moral principles, and relationships; e.g., Sutin & Robins 2007, Stawarczyk et al. 2011, Andrews-Hanna et al. 2013, Poerio et al. 2017; see also Johnson et al. 1988).

#### Exclusion Criteria and Quality Control

As is often observed in online experimentation, participant compliance varied. A series of quality control criteria were adopted to conservatively include only participants who fully engaged the task. The criteria were applied prior to any analysis of factors or assessments of reliability. Participants were excluded if they: 1) spent less than 20 min on the survey, 2) reported being outside of our age range (18-27 years-old), 3) had no mouse-clicks registered for multiple trial responses (indicating potential automation; Buchanan & Scofield 2018), or 4) showed clear stereotyped patterns of responding across trials. Participants were also flagged if they did not comment or write ‘None’ (as requested) in a final feedback box, or if they missed any check questions (i.e., they did not select “fully focused”, did not choose “cats and dogs” as popular pets or did not respond as “reading this question” on a question included specifically as a compliance check). A single flag (e.g., selecting “somewhat focused” or not writing ‘None’) did not result in exclusion if no patterned responding or other flags were noted.

Assessment of response patterns was particularly vital to quality control. 13 strategy probe ratings for 11 trials (7 that appeared across surveys and 4 unique to a survey) were visualized for each participant by two independent experimenters (LMD and OIA). Strategies were unlabeled during visualization to prevent experimenter bias. A participant was flagged for exclusion if strategy ratings did not differ across or within each trial, or if trials became uniform near the end of a survey. For example, a subset of participants chose a single value for all strategies within each trial (i.e., straight lines in the trial plots), a subset selected values in a clearly stereotyped pattern (e.g., 1-2-3-4-3-2-1-2-3-4 ratings across strategies), and a subset showed evidence of a drop off in performance (with a single rating given to all strategies, only in later trials). For participants flagged due to stereotyped responses, all trials were visualized as an additional check.

As a final quality control check, ratings across strategies for the 5 trials consistently included across all participants were visualized to explore cohort differences. Strategy ratings for each trial showed similar patterns. A linear model testing for rating differences found no significant cohort effects (*F*(6, 553)= 0.37, *p*=0.90), supporting that a trial’s strategy pattern represented stable properties, rather than idiosyncratic aspects of the rating group. As the replication experiment will reveal, these conservative procedures yielded highly stable estimates of strategy use across independent groups of participants.

136 total participants ages 18 to 25 remained (77.7%) after exclusion (Table 2). Included participants [mean age = 22.7 yr (SD = 1.9 yr), 49% identifying as female] had a mean completion time ranging from 38.2 - 56.7 min across the 7 cohorts (mean = 45.2 min).

**Table 2.**
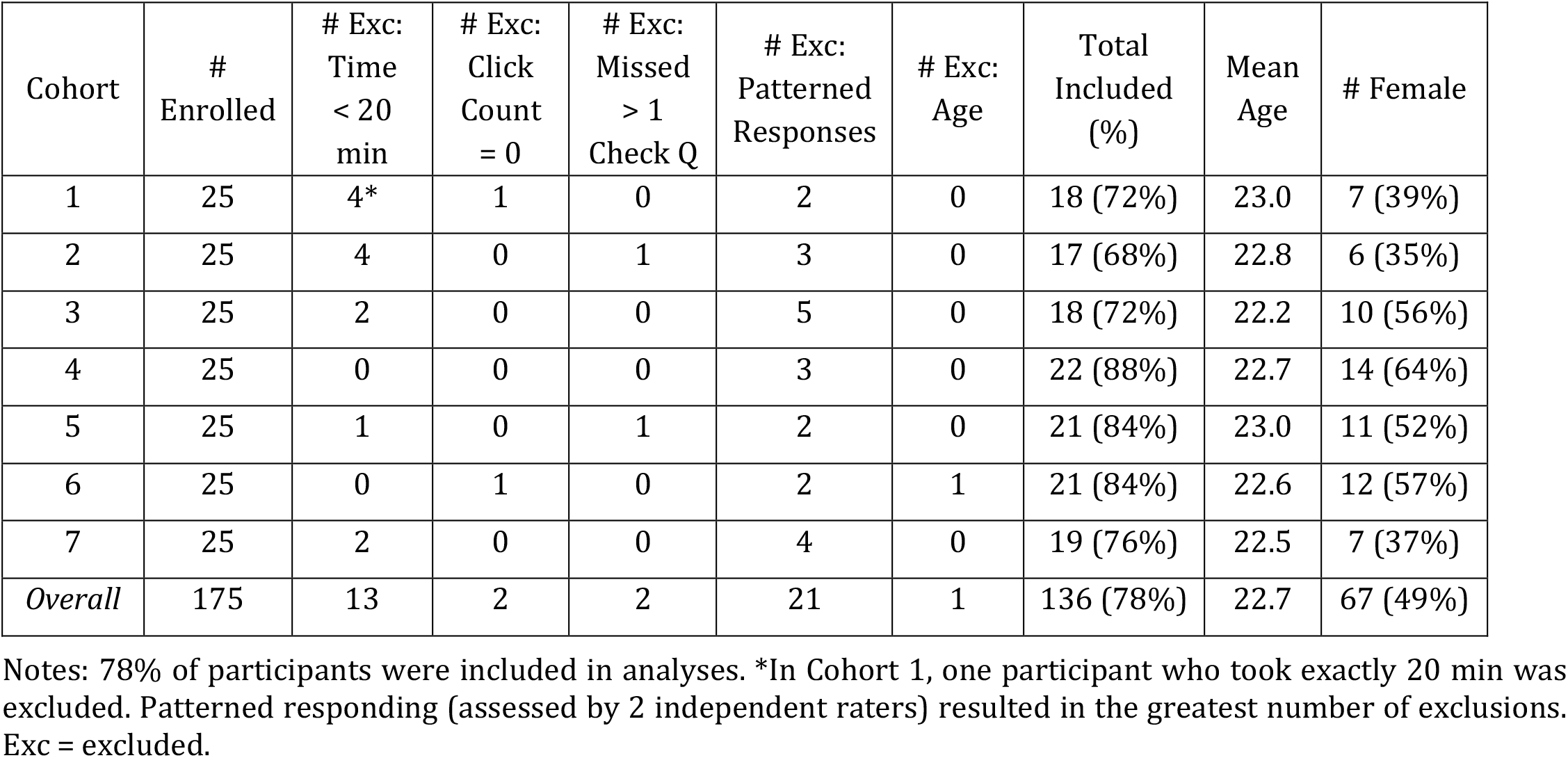
Summary of exclusions in Experiment 1.

| Cohort | # Enrolled | # Exc:<br>Time<br>< 20<br>min | # Exc:<br>Click<br>Count<br>= 0 | # Exc:<br>Missed<br>> 1<br>Check Q | # Exc:<br>Patterned<br>Responses | # Exc:<br>Age | Total<br>Included<br>(%) | Mean<br>Age | # Female |
| --- | --- | --- | --- | --- | --- | --- | --- | --- | --- |
| 1 | 25 | 4* | 1 | 0 | 2 | 0 | 18 (72%) | 23.0 | 7 (39%) |
| 2 | 25 | 4 | 0 | 1 | 3 | 0 | 17 (68%) | 22.8 | 6 (35%) |
| 3 | 25 | 2 | 0 | 0 | 5 | 0 | 18 (72%) | 22.2 | 10 (56%) |
| 4 | 25 | 0 | 0 | 0 | 3 | 0 | 22 (88%) | 22.7 | 14 (64%) |
| 5 | 25 | 1 | 0 | 1 | 2 | 0 | 21 (84%) | 23.0 | 11 (52%) |
| 6 | 25 | 0 | 1 | 0 | 2 | 1 | 21 (84%) | 22.6 | 12 (57%) |
| 7 | 25 | 2 | 0 | 0 | 4 | 0 | 19 (76%) | 22.5 | 7 (37%) |
| *Overall* | 175 | 13 | 2 | 2 | 21 | 1 | 136 (78%) | 22.7 | 67 (49%) |

Notes: 78% of participants were included in analyses. \*In Cohort 1, one participant who took exactly 20 min was excluded. Patterned responding (assessed by 2 independent raters) resulted in the greatest number of exclusions. Exc = excluded.

#### Strategy Clustering

Strategy clusters were identified with the goal of constructing robust composite scores for subsequent functional network analysis. We first calculated average ratings for each strategy, across respondents and for each unique trial. We *z*-scored ratings within strategies, creating a matrix featuring 16 mean strategy ratings x 180 trials. The raw correlation structure was estimated and visualized, and then hierarchical clustering was employed to estimate strategy groupings *(hclust* function and ward.D2 amalgamation procedure in R v3.5.1). We chose hierarchical clustering for the ease of visualizing relations across variables (see also Andrews-Hanna et al. 2013).

### Experiment 2: Prospective Replication of Strategy Rating Structure

Exp. 2 was conducted to examine the stability of online strategy ratings across trials, as well as to replicate the strategy clusters observed in Exp. 1 in an even larger sample, prior to network exploration in the neuroimaging data.

#### Participants

300 paid participants ages 18-27 were again recruited from Amazon Mechanical Turk using Cloud Research (Litman et al. 2017). Participants were English-speakers within the United States who had high ratings for completion of prior studies (90%+ approval rating with at least 100 prior tasks approved). Each participant provided informed consent through an online protocol approved by the Institutional Review Board (IRB) of Harvard University. Question and strategy probe forms were created and administered using Harvard’s Qualtrics platform.

#### Question and Strategy Probe Format, Exclusion Criteria and Quality Control

Exp. 2 followed the same procedures for data acquisition, QC and clustering as in Exp. 1, but with 300 total participants yielding 50 participants rating each unique trial question. Each participant received a subset of 38 total trial questions taken from the episodic projection task’s set of 180 trials. After QC exclusion, 238 participants ages 18 to 25 remained (79.3%) [mean age = 22.5 yr (SD = 1.9 yr), 61% identifying as female] with a completion time ranging from 38.1 - 46.6 min across the 6 cohorts (mean = 42.5 min). Analysis of strategy ratings across the same subset of 5 trials as in Exp. 1, repeated for all subjects, again revealed similar patterns and no cohort effect (*F*(5, 474=0.20, *p*=0.96).

#### Quantifying Behavioral Rating Stability Across Experiments

To examine the stability of measures derived from behavioral strategy probes, we compared the strategy patterns, for each trial, across independent groups of respondents from Exp. 1 and Exp. 2. For each trial, we plotted the mean (and standard error) across all 16 strategies from the RSS and calculated trial-level inter-experiment reliability (correlations across mean strategy ratings for that unique trial). Varying numbers of participants contributed to the mean ratings across trials, depending on the final participants retained after blind QC for compliance. For non-repeated trials, the number of respondents estimating strategy use for each unique trial was no fewer than 17 in Exp. 1 (mean = 19.4 across cohorts, max = 22; see Table 2) and no fewer than 37 in Exp. 2 (mean = 39.7 across cohorts, max = 42, see Table 3). Across experiments, 2-4 trials were repeated for subsets of cohorts (N_Exp.1_ = 72-117, N_EXP.2_ = 159 or 198), and at least 5 additional trials across all cohorts (N_EXP.1_ = 136, N_EXP.2_ = 238).

**Table 3.** Summary of participant exclusions in Experiment 2.

| Cohort | # Enrolled | # Exc:<br>Time<br>< 20<br>min | # Exc:<br>Click<br>Count =<br>0 | # Exc:<br>Missed<br>> 1<br>Check Q | # Exc:<br>Patterned<br>Responses | # Exc:<br>Age | Total<br>Included<br>(%) | Mean<br>Age | # Female |
| --- | --- | --- | --- | --- | --- | --- | --- | --- | --- |
| 1 | 50 | 3 | 1 | 1 | 8 | 0 | 37 (74%) | 22.9 | 21 (57%) |
| 2 | 50 | 5 | 0 | 0 | 6 | 0 | 39 (78%) | 22.3 | 26 (67%) |
| 3 | 50 | 5 | 0 | 1 | 3 | 1 | 40 (80%) | 22.4 | 18 (45%) |
| 4 | 50 | 4 | 0 | 1 | 3 | 0 | 42 (84%) | 22.5 | 25 (60%) |
| 5 | 50 | 2 | 1 | 0 | 6 | 0 | *40 (80%) | 22.6 | 27 (68%) |
| 6 | 50 | 3 | 0 | 1 | 4 | 2 | 40 (80%) | 22.3 | 29 (73%) |
| *Overall* | 300 | 22 | 2 | 4 | 30 | 3 | 238 (79%) | 22.5 | 146 (61%) |

Notes: 79% of participants were included in analyses. Patterned responding again resulted in the greatest number of exclusions. \*In cohort 5, one individual was excluded due to reported aphantasia (i.e., inability to represent visual imagery). Exc = excluded.

In addition, to quantify whether a strategy probe was rated similarly, *across* trials, from one experiment to the next, we calculated probe-level inter-experiment reliability. For each probe, we first calculated the mean ratings (and SD) across all 180 trials within each experiment. We then correlated the means across experiments (reported in Table 1).

#### Validating Composite Scores Against Trial Response Times

Strategy composite scores were based on self-reported strategy use. While reliability could be established for trials, probe questions and composites, a specific challenge of our approach is validation. As will be seen in the results, one behavioral composite emerged that reflected trial difficulty, creating an opportunity to test the validity of subjective ratings using response time (RT). For each trial, a mean RT was calculated, after exclusion of rare outlier trials with an RT greater than 60 sec (0.17% of all trials). These objective RT measures of trial difficulty were compared to the composite scores derived from the self-reported strategy use.

### Experiment 3: Examining Strategy-Network Relations

Exp. 3 examined trial-level variation in functional MRI responses to explore relations to trial-level strategy composite scores. The MRI data were included in a prior report by DiNicola et al. (2020) analyzed at the level of condition contrasts. Here the data were reanalyzed at the level of individual trials in the context of the novel behavioral data from Exp. 1 and Exp. 2.

#### Participants

Ten paid adult participants ages 18 to 25 [mean age = 20.5 yr (SD = 2.1), 9 right-handed, 8 identifying as female] were recruited from the Boston area to complete 4 neuroimaging sessions each. All participants provided informed consent through a protocol approved by the Harvard University IRB. Each neuroimaging session featured a battery of tasks, including fixation and the episodic projection task. Full details of task acquisition and preprocessing parameters are provided in DiNicola et al. 2020 (see also Braga et al. 2020). Only participants who completed all runs of the expanded episodic projection tasks were included in the current study (Exp. 2 and Exp. 3 in DiNicola et al. 2020). Two individuals who completed only two scanning sessions were excluded (S9 and S13 in DiNicola et al. 2020).

#### Fixation and Episodic Projection Task Paradigms

Each participant completed 11 runs of a passive fixation task, for intrinsic functional connectivity analysis (7 min 2s each, 77m total), as well as 6 runs of an episodic projection task (10 min 12s each, 61 min 12s total). Additional tasks were included in the neuroimaging battery, not discussed here (e.g., Braga et al. 2020, DiNicola et al. 2020).

During fixation, participants were instructed to fixate a black plus sign on a light grey background, while remaining alert and still. Fixation runs were intermixed with runs of other tasks, and fixation data were used for functional connectivity analysis to estimate network organization. Critically, networks were identified within each individual independently from (and prior to) other task analyses and without examination of any of the trial-variation effects explored in this paper.

During each run of the episodic projection task, participants responded to trials varying in self-relevance (Self vs. Non-Self) and temporal orientation (Past, Present, or Future). Crossing of these two dimensions (2 x 3) yielded 6 target conditions of 30 trials each (e.g., Past Self, Past Non-Self and so on for Present and Future timeframes; DiNicola et al. 2020, see also Andrews-Hanna et al. 2010). 5 trials from each condition were presented per task run. Across the 6 runs, each participant performed all 180 unique trials. Participants were instructed to carefully consider the details of each trial’s question before selecting a response (10s trial, 10s ISI).

#### MRI Data Acquisition and Processing

Data were acquired at the Harvard Center for Brain Science using a 3T Siemens Prisma-fit MRI scanner and a 64-channel phased-array head-neck coil (Siemens Healthcare, Erlangen, Germany). During each scan session, a rapid T1-weighted structural image was acquired, using a multi-echo magnetization prepared rapid acquisition gradient echo (ME-MPRAGE, van der Kouwe et al. 2008) sequence (1.2mm isotropic voxels, TR=2200ms, TE=1.57, 3.39, 5.21, 7.03ms, TI=1100ms, 176 slices, flip angle=7°, matrix=192 × 192 × 176, in-plane GRAPPA acceleration=4). Blood oxygenation level-dependent (BOLD) data were acquired using a multi-band gradient-echo echo-planar pulse sequence (see Feinberg et al. 2010, Moeller et al. 2010, Setsompop et al. 2012, Xu et al. 2013), provided by the Center for Magnetic Resonance Research at the University of Minnesota (2.4mm isotropic voxels, TR=1000ms, TE=32.6ms, flip-angle=64°, matrix=88 × 88, 65 slices covering cerebral cortex and cerebellum). All data were processed using a custom analysis pipeline (termed ‘iProc’; see Braga et al. 2019, DiNicola et al. 2020), designed to preserve within-individual details. Briefly, each participant’s data were registered to a subject-specific, 1mm-isotropic T1 template through a single interpolation, which combined matrices for motion, field map unwarping, alignment to a mean BOLD image and then to the T1 template.

For functional connectivity analysis, nuisance variables (6 motion parameters and whole-brain, ventricular and white matter signals, along with their temporal derivatives) were regressed from the T1-aligned BOLD fixation data. These data were then bandpass filtered at 0.01-0.10 Hz (using AFNI v2016.09.04.1341; Cox 1996, 2012). For episodic projection task analyses, the whole brain signal was regressed from the T1-aligned task data (see DiNicola et al. 2020). All BOLD data from both the fixation task and episodic projection task were then resampled to the fsaverage6 cortical surface mesh (using trilinear interpolation; Fischl et al. 1999) and smoothed using a 2mm full-width at half-maximum kernel.

Data were examined for quality. Run-level exclusion criteria included: (1) maximum absolute motion greater than 1.8mm, (2) signal-to-noise ratio less than or equal to 135, and, for the episodic projection tasks, (3) eyes closed during skipped task trials. 16 fixation runs were excluded across the participants. No episodic projection task runs were excluded (see DiNicola et al. 2020 for behavioral performance).

Functional connectivity (FC) analyses were conducted on fixation data, within each individual, in order to precisely estimate whole-brain network organization. *k-*means estimates of networks were used, as reported in Braga et al. 2020. Briefly, for each individual, medial wall vertices were removed, and then time series data from the fixation runs were *z*-normalized, concatenated and input to the *k*-means algorithm, using default parameters (MATLAB v2015b). Networks were identified within the whole-brain *k*-means outputs based on referential features (e.g., Braga & Buckner 2017). Parcellations were computed while varying *k* from 10 to 20, and the solution featuring the fewest clusters differentiating 6 networks was chosen for each individual (Braga et al. 2020). These networks included default network A (DN-A) and B (DN-B), a language network (LANG), frontoparietal control network A (FPN-A) and B (FPN-B), and a salience network (SAL). For 2 individuals, features of one network were observed in two clusters, which collectively better matched seed-based *post hoc* checks of the networks. Both were included in the network estimate (FPN-A for S10 and FPN-B for S3), prior to task analysis (Braga et al. 2020). The networks were determined fully before the task functional MRI data were examined to avoid any potential bias.

For the episodic projection task, run-specific GLMs were created featuring separate regressors for every trial (e.g., Hassabis et al. 2014), producing trial-specific beta-maps (see full details in DiNicola et al. 2020; GLMs created using FSL v5.0.4). Beta values within each network were then extracted and averaged, yielding trial-specific responses for each separate network within each participant (180 total trials).

Since the current analyses aimed to explore trial-level variation (i.e., differences in stable trial-level properties), after estimating average values for each network within individuals, we averaged trial values across individuals, producing a single network estimate for every unique trial. In this way, within-individual network definition allowed for estimates of network activity that were fully constructed within the idiosyncratic anatomy of each individual. At the same time, extremely stable functional MRI estimates for each of the 180 individual trials were obtained because each trial estimate was the average across the 10 participant’s individualized networks.

#### Examining Strategy-Network Relations

Each of the 180 trials from our episodic projection task was paired with 5 mean strategy composite scores (from the online data) and 6 mean network activity values (from the neuroimaging data). We asked whether variation in our behavioral composite scores, across trials, related to variation in network activity. As a first step, we calculated correlations between trial-level behavioral composite scores and mean functional MRI BOLD response estimates from each of the 6 networks. Data from all trials were included, and Pearson’s correlation values were plotted to visualize patterns for each composite.

As will be seen in the results, particularly strong correlations were found between the Scene Construction composite scores and DN-A response and between the Difficulty composite scores and FPN-B response. Building on these correlational results, we next sought to unpack observed strategy-network relations. Scene Construction, for example, appeared to differentiate DN-A from tightly juxtaposed DN-B, as well as other networks. Relations between Scene Construction and DN-A and Difficulty and FPN-B appeared dissociable. We sought to test these observations using multiple methods.

To quantify whether composite scores significantly predicted network activity, multiple regression was employed. Each regression model probed whether composite scores explained variance in a specific network’s response. The relative importance (i.e., R^2^ contribution of each regressor) for each composite was also calculated (*relaimpo* R; Grömping 2006). Scatterplots allowed for visualization of different composite-network relations, relative to regression lines and across all 180 trials.

Regression models thus provided insight into how much variance in network activity was captured by our composite scores. For the strongest observed correlations, we also sought to contextualize these values as a percent of the *explainable* variance - the highest R^2^ one could expect, constrained by the internal reliability of our data (i.e., given a true correlation of 1 between a network-composite pair; e.g., Konkle et al. 2010; see Vul et al. 2009). Toward this goal, we first calculated split-half reliability for the composites and networks, by halving each dataset, creating vectors of mean values for each half, and correlating those values (adjusted by the Spearman-Brown formula). For network estimates, all split-half combinations were used, and for composite estimates, 1,000 split-half samples (featuring 36 participants per strategy probe from Exp. 2). A maximum explainable variance value was estimated as the product of the reliability, which we compared to our estimated R^2^ values.

#### Exploring the Impact of Difficulty

As will emerge in the results, initial regression models revealed evidence of inter-correlation between a composite score reflecting Difficulty and the other composite scores. In *post hoc* analyses we directly tested whether Difficulty impacted other composite-network relations by regressing the impact of Difficulty ratings from all other composite scores and plotting residual composite-network correlations.

#### Probing Scene Construction and Difficulty Contrasts to Verify Network Dissociations

Finally, the analyses supported a double-dissociation between composite scores reflecting Scene Construction and Difficulty in relation to activity in brain networks DN-A and FPN-B. As a final stringent test of this discovery, we used the composite scores to create contrast maps based on trials with high and low values on each of these composites. For Difficulty, we identified the 10 trials with the highest and 10 trials with the lowest composite scores. For each individual, we then created a whole-brain contrast map, and overlaid the border of the individual’s FPN-B, to compare each individual’s contrast map to their specific FPN-B estimate. For Scene Construction, we used the same process, but *only* included trials originally considered controls (i.e., from Present Self, Past Non-Self and Future Non-Self conditions). We selected the 10 controls trials with the highest and 10 controls trials with the lowest Scene Construction composite scores, and then visualized resultant contrast maps in relation to each individual’s DN-A estimate.

## Results

### Behavioral Strategy Probe Ratings Capture Stable Trial-to-Trial Variance

Nearly every strategy probe showed high inter-experiment reliability (all *r* > 0.80 except Specificity; see Table 1). Within each trial, mean ratings from the independent cohorts were also strikingly similar (mean *r* = 0.94 across trials, *r* > 0.80 for 98% of trials; see Figures 1 and 2 for examples). These stable patterns across raters provided evidence that trial ‘traits’ could also inform brain network activity (as indirectly estimated by BOLD functional MRI) from the independent neuroimaging sample.

In addition, trials showed trial-to-trial variation, including within originally-designed task conditions. Trials designed to target episodic projection, for example, were previously shown - on average - to preferentially recruit DN-A (DiNicola et al. 2020). But strategy patterns varied substantially among individual episodic projection trials (see Figure 1), raising the question of *which* component dimensions might explain DN-A recruitment. Original control trials (i.e., designed *not* to require episodic memory or prospection) also showed marked variability (see Figure 2). The variation went well beyond differences between conditions. For example, while on average, control trials exhibited lower reliance on the personal past than target trials, as intended, multiple control trials also showed high ratings on strategies relevant to mental scenes or events, also observed in target trials’ patterns. These results highlighted the opportunity to leverage trial-level variation toward novel exploration of network processes, beyond condition-level distinctions.

### Five Strategy Composite Scores are Supported by Hierarchical Clustering

To visualize relations among the 16 strategies, we created a correlation matrix, using data from all 180 trials. In Exp. 1, the matrix revealed strong correlations between groups of strategies (Figure 3, left), adding to evidence that trials feature distinct rating combinations. Trials with high ratings for visual imagery (“Visual_Imagery”), for example, were also likely to have high ratings for envisioning the physical locations of objects and places (“Loc_Obj_Places”). Trials with high ratings for considering others’ mental states (“Others_Feelings”) tended also to require imagining others’ personality traits (“Others_Personality”). In Exp. 2, the correlation matrix showed similar structure (Figure 3, right), supporting the reliability of inter-strategy correlations.

**Figure 3.**
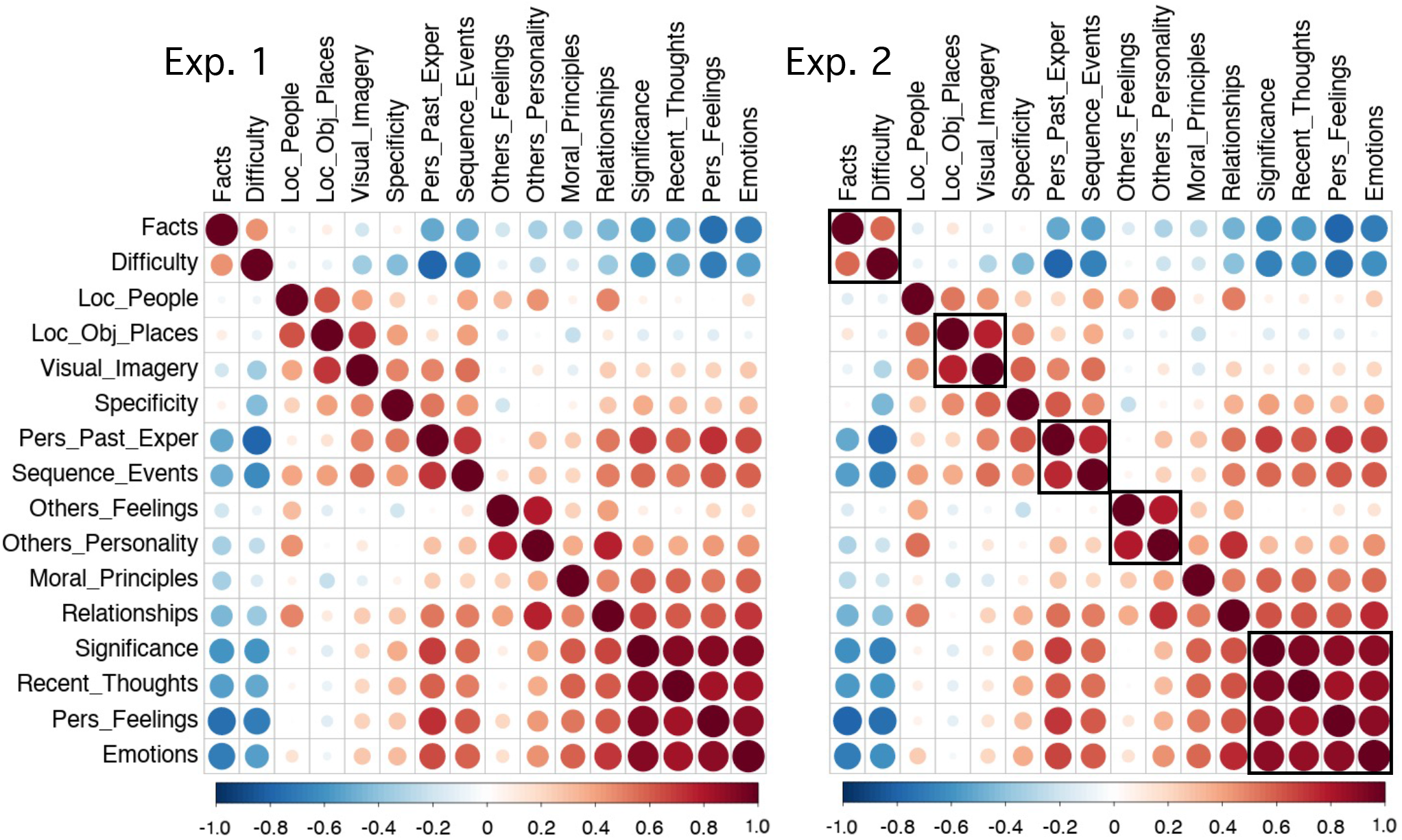
Reliable strategy clusters emerge that capture trial-to-trial variation. **(Left)** A correlation matrix illustrates the relations among the 16 scaled strategy probes, using data from all 180 trial questions in behavioral Exp. 1. Strong correlations emerge between subsets of strategy probes indicating that individual trials have distinct rating combinations. For example, trials high in use of visual imagery (Visual_Imagery) also tend to be high in reports of imagining the locations of objects and places (Loc_Obj_Places) and, to a lesser degree, the locations of people (Loc_People). **(Right)** An independent correlation matrix from Exp. 2 reveals that the strategy relations are reliable (ordered here as in Exp. 1 for visualization). The boxes around correlation clusters in Exp. 2 reveal the groupings that were selected based on hierarchical clustering as shown in Figure 4. Pers = Personal; Exper = Experiences.

Using hierarchical clustering, we next identified 5 strategy groupings that could be combined into composite scores for subsequent network analysis. In Exp. 1, the greatest strategy differentiation appeared for a pair of correlated strategies (“Facts” and “Difficulty”; see Figure 3, left). We cut the clustering dendrogram to preserve clusters at least as strong as this pair, which revealed 5 groupings (Figure 4, top). In Exp. 2, the independently estimated dendrogram revealed a similar structure. Using the same cut point (above “Facts” and “Difficulty”) produced 5 strategy groupings that largely matched those from Exp. 1, which we heuristically labeled: (I) Difficulty, (II) Autobiographical, (III) Scene Construction, (IV) Others-Relevant, and (V) Self-Relevant.

**Figure 4.**
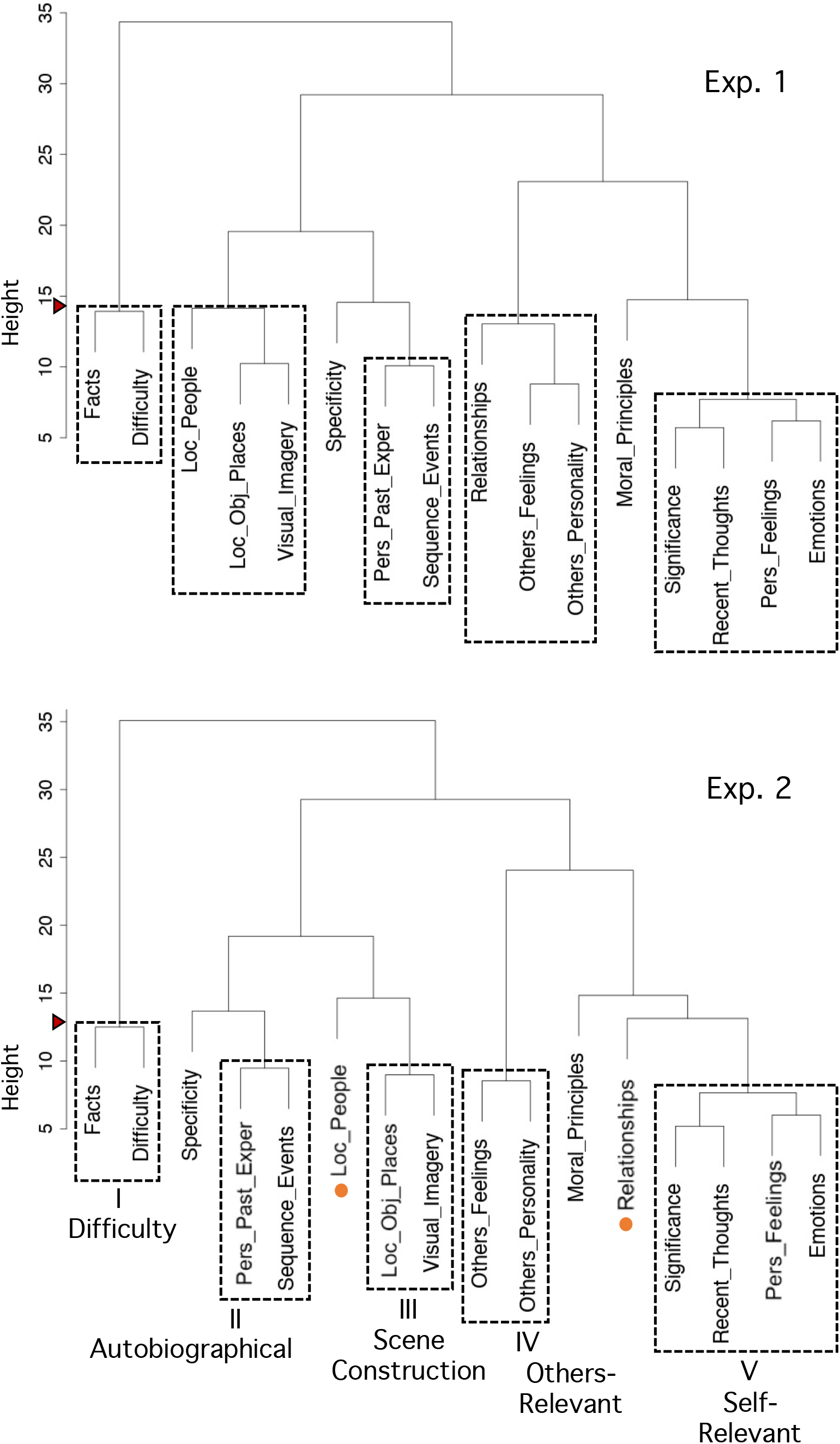
Hierarchical clustering yields five distinct strategy composite scores. Hierarchical clustering identified strategies that could be combined into composite scores for functional network analysis. **(Top)** The dendrogram from Exp. 1 displays composites using a cut point above “Facts” and “Difficulty,” preserving all clusters at least as strong as this pair. The cut point is noted by a red triangle on the y-axis. Dashed boxes show the cluster groupings. **(Bottom)** The independently estimated dendrogram from Exp. 2 reveals a similar structure. Using the same cut point above “Facts” and “Difficulty” leads to similar clusters that include a core set of strategy probes converged upon across both experiments. These 5 strategy composites were carried forward for analysis of the functional MRI data. They are heuristically labeled as: (I) Difficulty, (II) Autobiographical, (III) Scene Construction, (IV) Others-Relevant, and (V) Self-Relevant. Strategy probes that were not consistent between the two experiments (or weakly associated) were not included in the final composite scores, allowing only the most robust and stable strategy probes to be incorporated into the final 5 composite scores.

The two strategies that were not identically grouped between Exp. 1 and Exp 2. (“Loc_People” and “Relationships”), as well as those that showed weaker grouping across both experiments (“Specificity”, “Moral_Principles”), were excluded from composite scores. Out of all strategy probes, “Specificity” also had the lowest inter-experiment reliability (*r* = 0.66) and “Moral_Principles” had the lowest mean rating and SD (likely reflecting few morally-relevant questions in our task trials; see Table 1).

Given that strategy ratings from Exp. 1 and Exp. 2 exhibited comparable correlational structure and clustering results, ratings from the larger dataset (Exp. 2) were used to create the strategy composite scores carried forward to analyses of the functional MRI data.

### Difficulty Composite Scores Correlate with Response Times Across Trials

Difficulty composite scores tracked RT values (Figure 5; *r* = 0.65; CI[0.56, 0.73]). Overall, results from Exp. 1 and 2 provided evidence that trial-level ratings were stable (Figures 1 and 2), captured trial-to-trial variation (Figures 1 and 2), and clustered in reliable ways across experiments (Figures 3 and 4). For Difficulty (the only possible instance), the composite score was validated against a separate objective measure (Figure 5). These findings all supported proceeding with analyses of the functional MRI data in relation to strategy composite scores.

**Figure 5.**
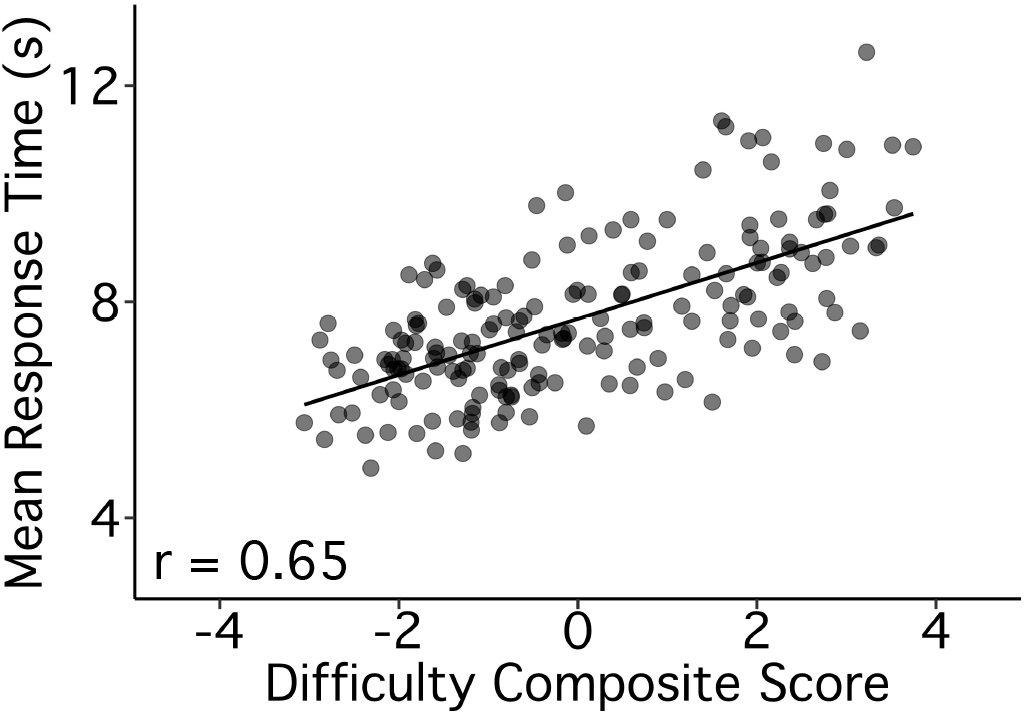
Difficulty composite scores track trial-to-trial variation in response times supporting validity. Response time (RT) estimates provided an opportunity to validate subjective ratings of Difficulty. Mean RTs were calculated for each trial (y-axis) and plotted against the Difficulty composite scores (x-axis) from Exp. 2. The observed strong positive relation provides evidence for the validity of the Difficulty composite, even though it is based on participant self-report. The Pearson’s correlation value is shown in the bottom left. The line represents a linear model predicting Difficulty scores by RT across trials.

### Strategy Composites Scores Correlate Differentially with Network Activity

We first calculated correlations between each of the 5 strategy composites and functional MRI response in each of the 6 independently-estimated networks (Figures 6 and 7), using data from all 180 trials. Plotting Pearson’s correlation values revealed a particularly striking relation between Scene Construction scores and DN-A activity (Figure 8). Scene Construction differentiated DN-A from interwoven DN-B and from all four additional networks. DN-B, in turn, showed a selective (albeit weaker) correlation to Others-Relevant scores. In addition, strong correlations were noted between the Difficulty composite score and both FPN-A and FPN-B. More ambiguous network-composite results were revealed for Autobiographical and Self-Relevant scores (due, in part, to confounding effects of Difficulty; see Figure 12).

**Figure 6.**
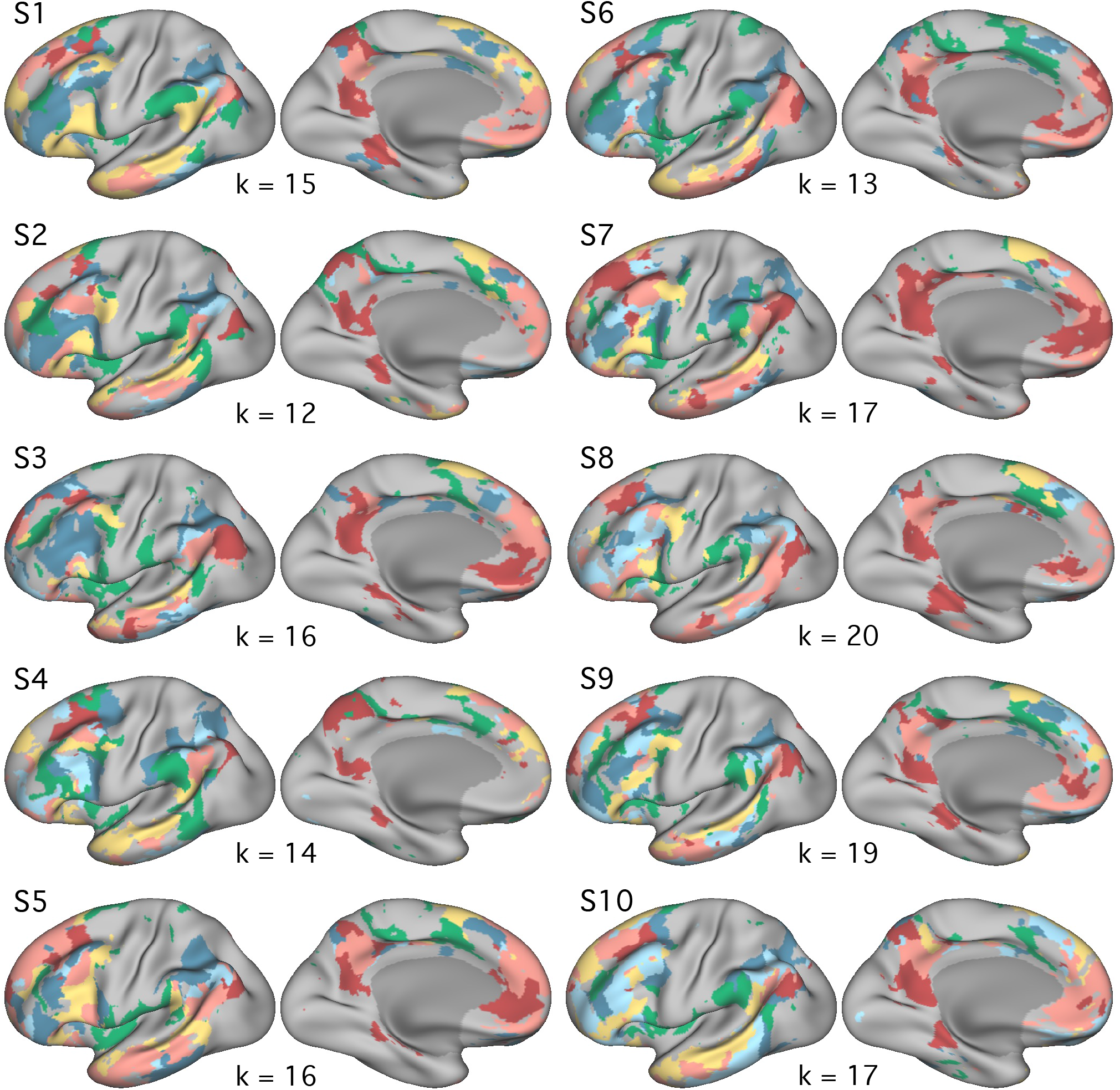
Distributed networks estimated from functional connectivity within individuals: Left hemisphere. Whole-brain estimates of the 6 networks are displayed for each of the 10 extensively-sampled individuals (identical to Braga et al. 2020; see also DiNicola et al. 2020). For each individual, the k-means solution featuring the fewest clusters that differentiated the 6 target networks was chosen. Networks are shown in the left hemisphere and include Default Network A (DN-A, red), Default Network B (DN-B, pink), a candidate Language Network (LANG, yellow), two candidate Frontoparietal Control Networks (FPN-A, light blue; FPN-B, dark blue) and a candidate for the Salience Network (SAL, green). These networks were defined independently of assessments of functional response properties.

**Figure 7.**
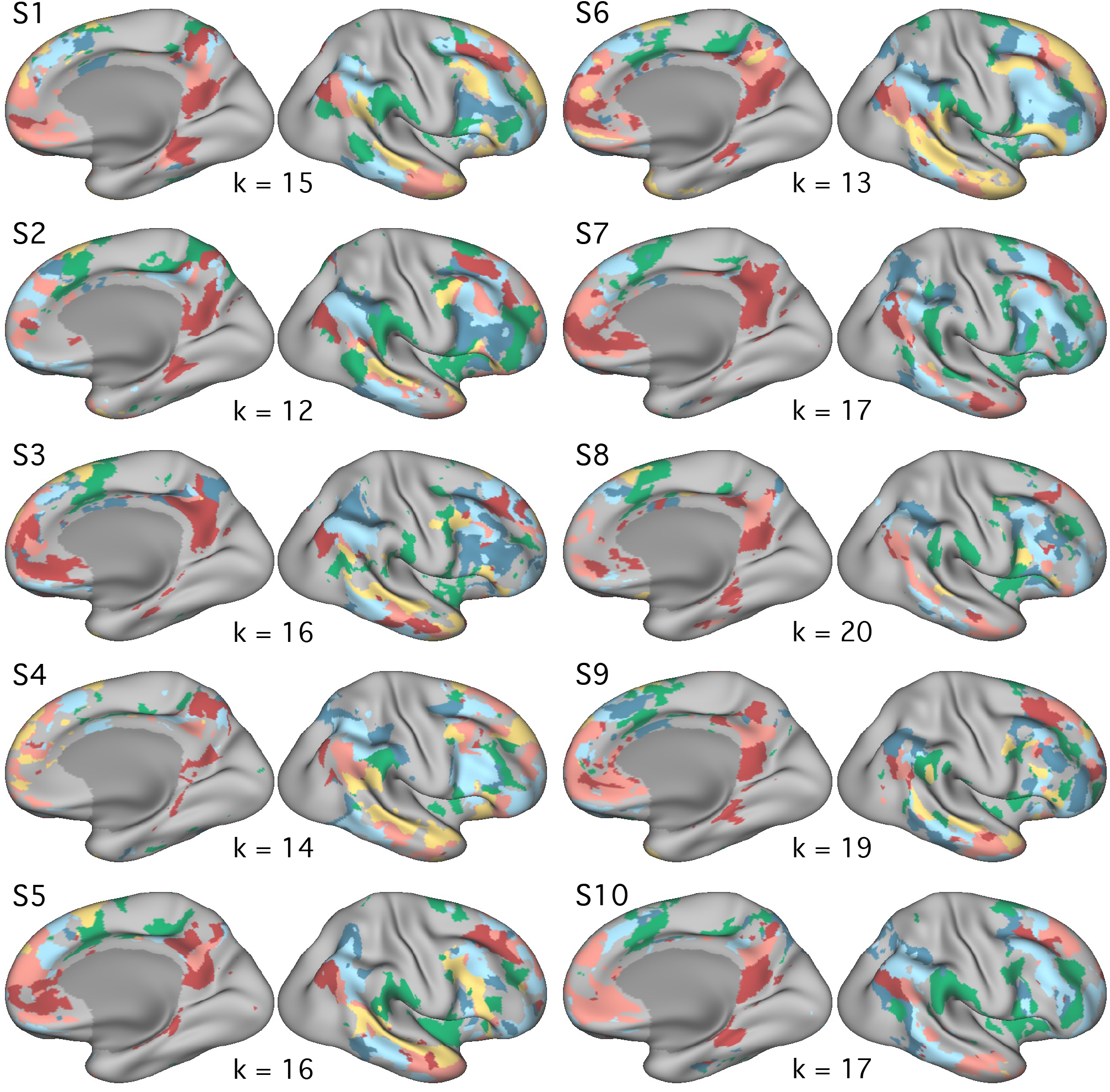
Distributed networks estimated from functional connectivity within individuals: Right hemisphere. The networks from Figure 4 are displayed for the right hemisphere, including Default Network A (DN-A, red), Default Network B (DN-B, pink), a candidate Language Network (LANG, yellow), two candidate Frontoparietal Control Networks (FPN-A, light blue; FPN-B, dark blue) and a candidate for the Salience Network (SAL, green).

**Figure 8.**
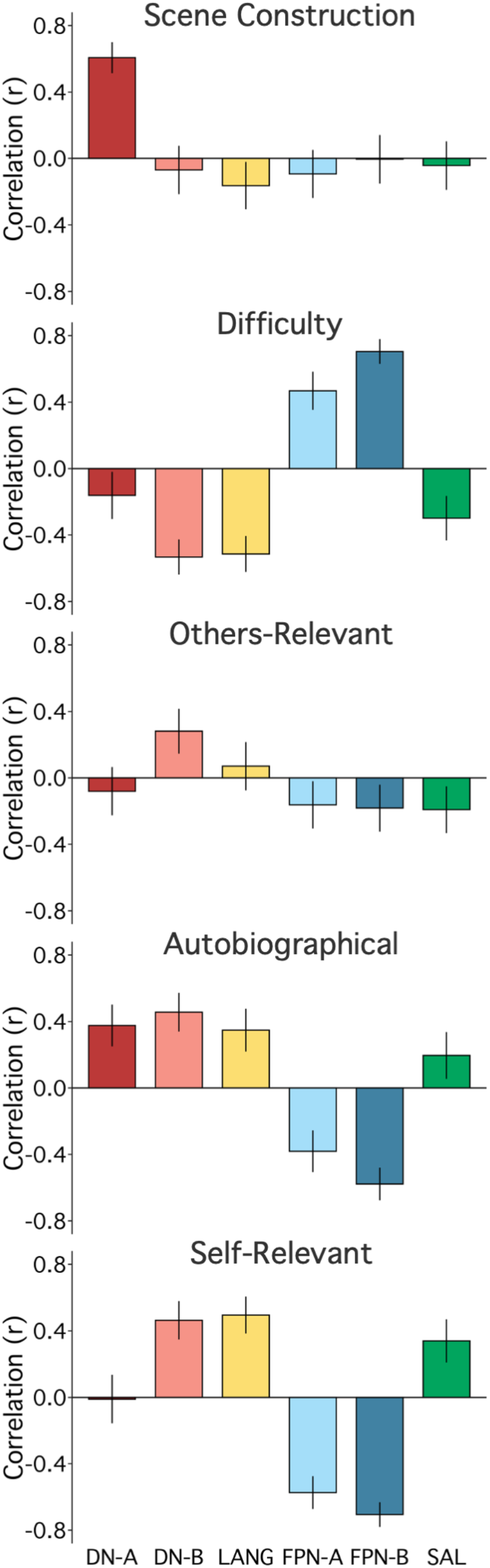
Strategy composites are associated with differential and selective network activity. For each strategy composite, scores from all 180 trials were correlated with the response estimates for each of the 6 networks. The strategy is labelled at the top of each plot; the six colored bars reflect the Pearson’s correlations with 95% confidence intervals. **(Scene Construction)** A particularly striking and selective relation is observed between Scene Construction composite scores and DN-A activity. **(Difficulty)** The Difficulty composite scores show a strong positive correlation to FPN-B and a strong but weaker relation to FPN-A. **(Others-Relevant)** The Others-Relevant composite scores reveal a modest association with DN-B. **(Autobiographical and Self-Relevant)** Results between the Autobiographical and Self-Relevant composite scores are more ambiguous (partially related to confounding effects of Difficulty; see Figure 12). Networks, from left to right: Default Network A (DN-A, red), Default Network B (DN-B, pink), a candidate Language Network (LANG, yellow), two candidate Frontoparietal Control Networks (FPN-A, light blue; FPN-B, dark blue) and a candidate for the Salience Network (SAL, green).

To unpack these findings, we first probed Scene Construction’s relation to DN-A and DN-B. The episodic projection task was originally designed to differentiate these parallel networks, using condition-level contrasts (DiNicola et al. 2020). The observed correlation suggested that DN-A might preferentially support Scene Construction processes, which would inform differentiation between these tightly juxtaposed networks (e.g., Buckner & DiNicola 2019, DiNicola et al. 2020, see also Deen et al. 2020).

### Scene Construction is Selectively Related to DN-A But Not DN-B Activity

Comparing Scene Construction scores, across trials, to DN-A activity revealed a strong positive correlation (Figure 9; *r* = 0.61, CI [0.51, 0.69]). Conversely, DN-B showed a weakly negative correlation to Scene Construction (Figure 9; *r* = −0.07, CI [-0.21, 0.08]).

**Figure 9.**
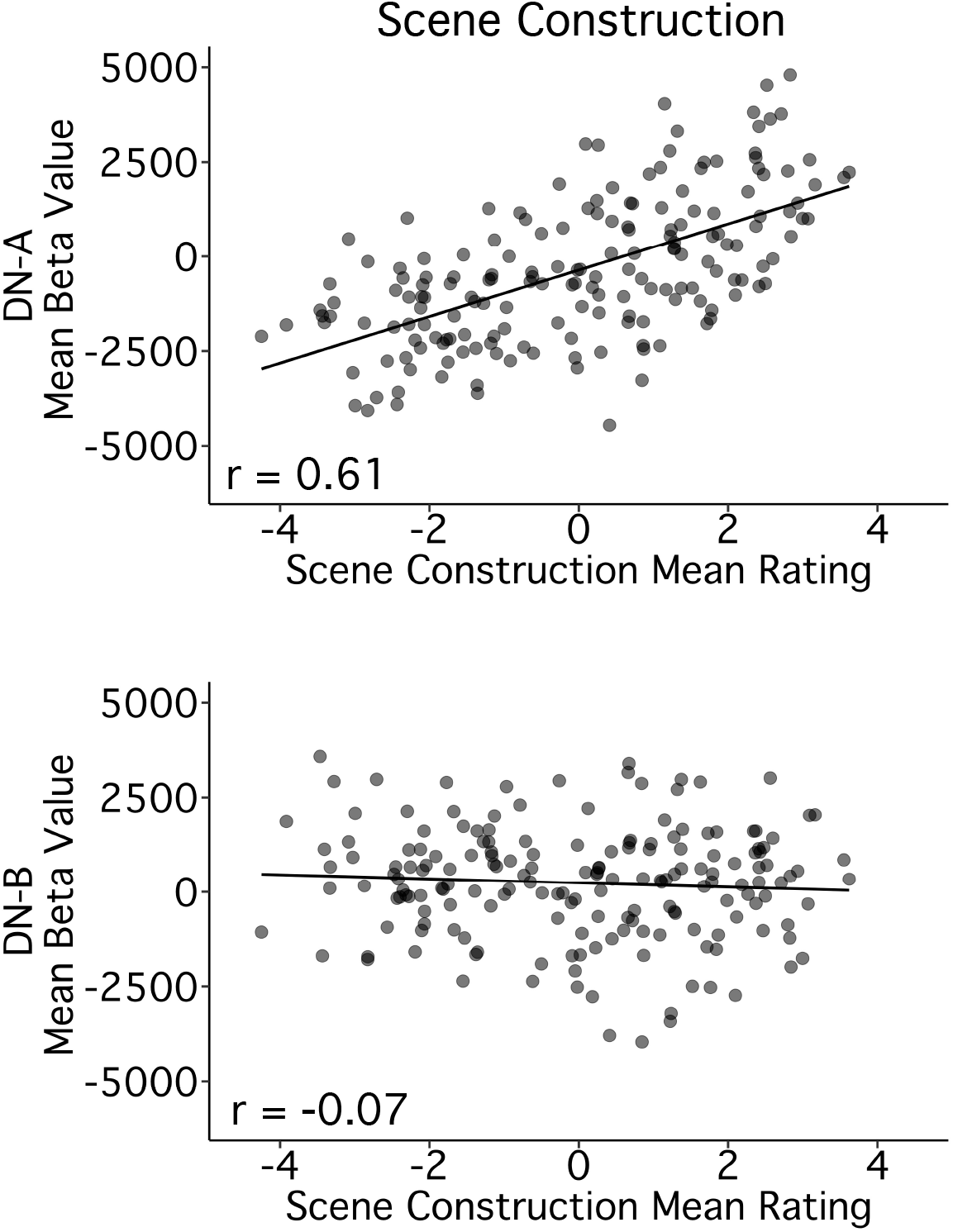
Scene Construction is selectively related to DN-A but not DN-B activity. Scatter plots of individual trial activity levels within DN-A illustrate a strong relation to the Scene Construction composite. **(Top)** The mean activity level for each of the 180 trials is plotted for DN-A (y-axis) against the mean Scene Construction composite scores (x-axis). Note that each separate point represents the mean behavioral score for that unique trial from 37-42 participants and the mean functional MRI response for that unique trial averaged across 10 participants. There is a striking linear relationship between the Scene Construction composite and DN-A activity. **(Bottom)** The mean activity level for each trial is similarly plotted for the adjacent network DN-B against the same Scene Construction composite scores (x-axis). There is minimal relation. Pearson’s correlation values are shown in the bottom left corners.

Multiple regression, with network-specific models, supported the observed dissociation. Of note, initial models included all 5 strategy composites, but variance inflation factors (VIF) revealed moderate intercorrelation between Difficulty, Autobiographical and Self-Relevant scores (VIF > 3; e.g., see Johnston et al. 2018). To avoid multicollinearity from a Difficulty confound (which can reduce model accuracy and introduce redundancy), we removed Autobiographical and Self-Relevant composites from subsequent models (see also Figure 12). Remaining VIF factors suggested no persisting multicollinearity (i.e., equaled 1), so we interpreted regression models featuring Difficulty, Scene Construction and Others-Relevant composite scores.

For DN-A, this model significantly predicted activity (*F*(3,176) = 36.91, *p* < 0.001), and Scene Construction was the only significant predictor, accounting for most of the variance in DN-A response (R^2^_Scene Construction_ = 0.36, *p* < 0.001; R^2^_Full Model_ = 0.39). For DN-B, the overall model was significant (*F*(3,176) = 28.55, *p* < 0.001), but Scene Construction was not a significant predictor and accounted for almost no variance in DN-B response (R^2^_Scene Construction_ = 0.01, *p* > 0.05; R^2^_Full Model_ = 0.33). These results supported a selective relation between Scene Construction scores and DN-A activity. Although DN-B is tightly juxtaposed to DN-A across the cortical mantle (e.g., Braga & Buckner 2017), DN-A appears to play a distinct role in Scene Construction processes.

As an additional consideration, the Scene Construction composite included strategies for using visual imagery and for considering the locations of objects or places. A third strategy, for considering the locations of people, clustered with this composite in Exp. 1 and was related, but more weakly, in Exp. 2. To test whether excluding this strategy impacted the observed dissociation between DN-A and DN-B, in *post hoc* analyses, we created a separate composite that included “Visual_Imagery”, “Loc_Obj_Places”, and “Loc_People” ratings. Findings were comparable. For DN-A, this composite (replacing Scene Construction in the original model) was still a significant predictor, again accounting for most of the variance in DN-A response (R^2^ = 0.38, *p* < 0.001; R^2^_Full Model_ = 0.43; all three predictors now *p* <0.05). For DN-B, this composite was not a significant predictor (R^2^ = 0.00, *p* > 0.05; R^2^_Full Model_ = 0.32).

### Trial-Level Variation in Scene Construction Tracks DN-A Activity, Including for Control Trials

Given that the episodic projection task was designed to target DN-A activity (DiNicola et al. 2020), we next aimed to test whether a link between Scene Construction and DN-A simply recapitulated previous, condition-level results. Namely, in prior work, trials from the Past and Future Self conditions were shown to preferentially recruit DN-A (DiNicola et al. 2020). Correlations between DN-A response and Scene Construction scores might predominantly reflect the same condition-level distinctions.

We therefore ran additional analyses restricted either to those original target trials *or* to control trials constructed *not* to include episodic projection demands (from Present Self, Past Non-Self and Future Non-Self conditions). Correlation results revealed that Scene Construction scores for *both* the original target (*r* = 0.33, CI[0.19, 0.45]) and original control trials (*r* = 0.48, CI[0.35, 0.58]) tracked DN-A response (Figure 10). Although Scene Construction scores were higher, on average, for the target trials as compared to the control trials (t(146) = −10.82, *p* < 0.001; see triangles in Figure 10), Scene Construction ratings were significant predictors of DN-A activity in models restricted to the control trials and still accounted for the most variance (R^2^_Scene Construction_ = 0.19, *p* < 0.001; R^2^_Full Model_ = 0.32, Difficulty also *p* < 0.01).

**Figure 10.**
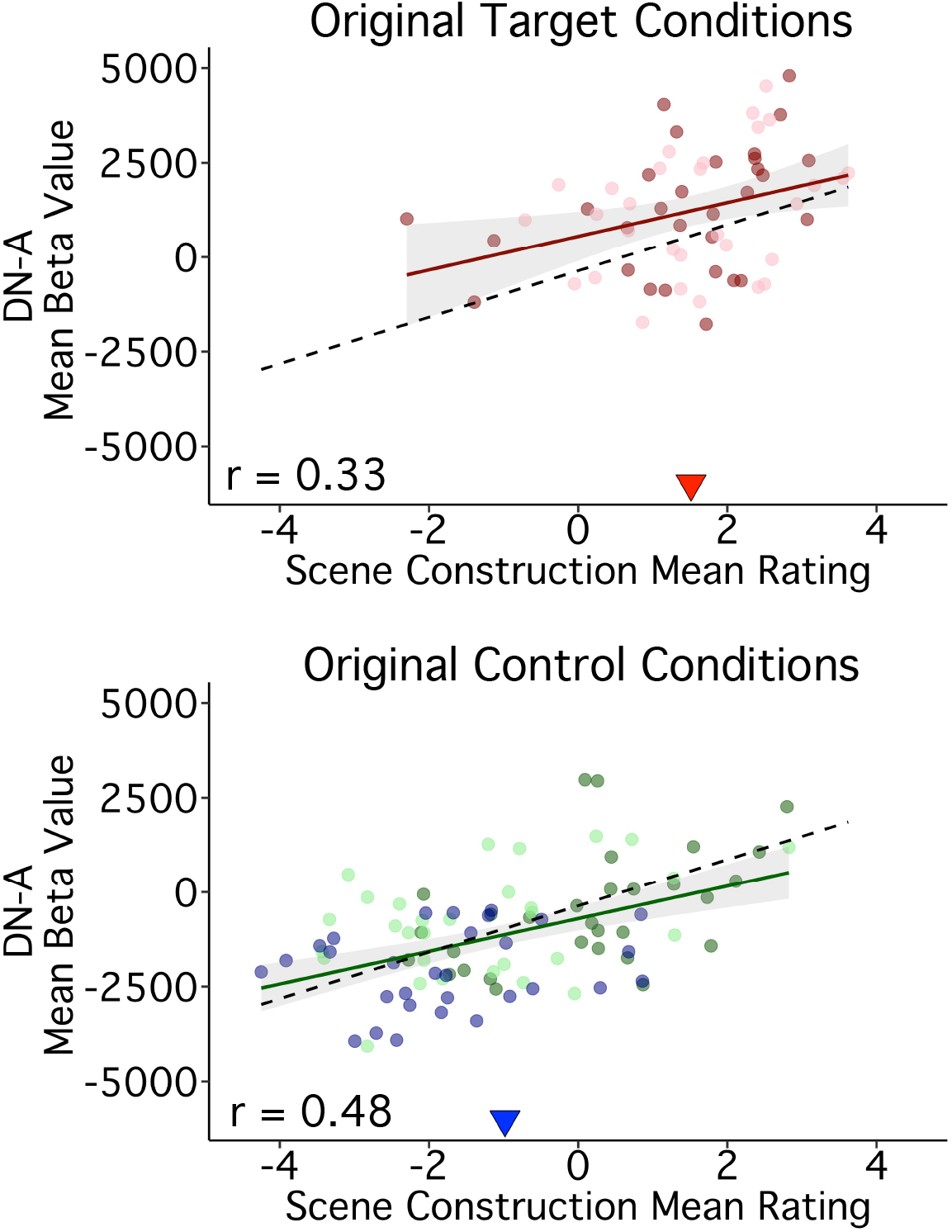
Trial-to-trial variation in Scene Construction tracks DN-A activity levels even for trials that do not involve episodic remembering or prospection. Scatter plots are displayed for DN-A split by whether the trials originated from the target conditions constructed to demand episodic projection **(Top)** or were originally included within control conditions constructed to minimize such demands **(Bottom)**. The correlation with Scene Construction is present in both sets of trials with a strong and clear linear association within the original control trial conditions. Points are colored based on their condition origin: Past Self (pink), Future Self (red), Present Self (blue), Past Non-Self (dark green) and Future Non-Self (light green). Each point represents a single trial; Pearson’s correlation values are shown in the bottom left corners. Regression lines in color include only the trials from target **(Top)** or control **(Bottom)** conditions; black dashed regression lines include all 180 trials. Triangles indicate the mean composite score across trials in each plot, highlighting condition-level differences. Scene Construction tracks DN-A response *both* for trials originally constructed to target DN-A and for trials constructed to minimize demands on episodic projection.

To reiterate, even for trials designed *not* to require episodic projection, the extent to which trials required scene construction predicted DN-A activity. This provides evidence that a core process subserved by DN-A, regardless of episodic projection, is mental construction of scenes (see also Hassabis & Maguire 2007, Hassabis & Maguire 2009).

### Scene Construction and Difficulty Support a Robust Functional Double Dissociation Between DN-A and Another Juxtaposed Network FPN-B

To further examine network heterogeneity, we leveraged the composites with the strongest network correlations - Scene Construction and Difficulty - toward probing potential functional dissociation between DN-A and FPN-B. These networks also feature juxtaposed regions in multiple cortical zones (Figures 6 and 7; see also Braga & Buckner 2017, Braga et al. 2020).

As described, Scene Construction scores correlated to DN-A activity (*r* = 0.61, CI [0.51, 0.69]). Conversely, DN-A showed a weakly negative correlation to Difficulty scores (r = −0.16, CI [-0.30, −0.02]), which were strongly associated with activity in the adjacent FPN-B (*r* = 0.70, CI [0.62, 0.77]). FPN-B, in turn showed no relation to Scene Construction (*r*= 0.00, CI [-0.15, 0.14]), illustrating a functional double dissociation (Figure 11).

**Figure 11.**
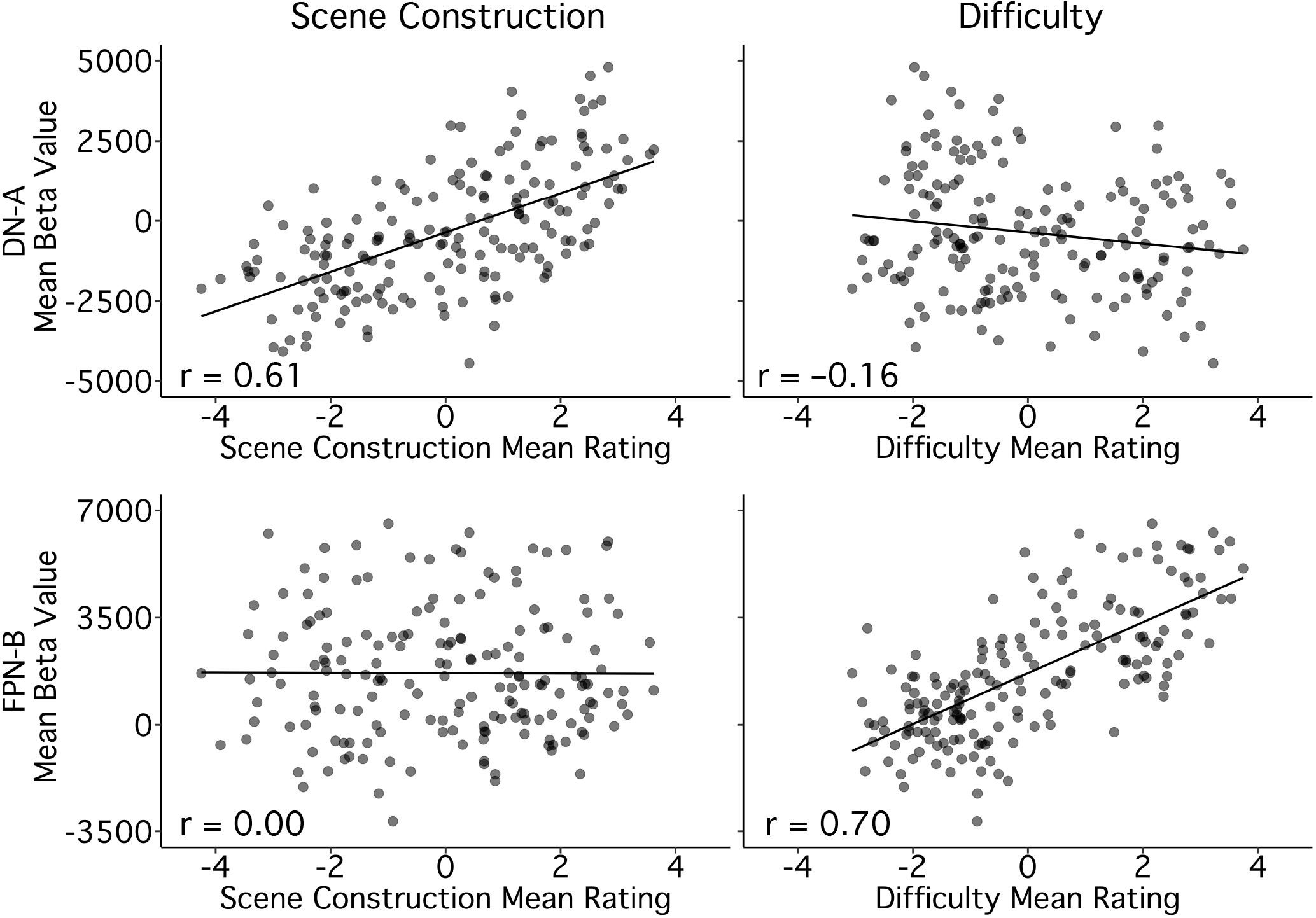
Contrasting Scene Construction and Difficulty reveals a double-dissociation between DN-A and FPN-B. Scatter plots contrast the differential relations of DN-A and FPN-B with the Scene Construction and Difficulty composite scores. **(Top, Left)** The mean activity level for each of the 180 trials is plotted for DN-A (y-axis) against the mean Scene Construction composite scores (x-axis). **(Bottom, Left)** The mean activity level for each trial is plotted for FPN-B (y-axis) against the mean Scene Construction composite scores (x-axis). Note the absence of a relation. **(Top, Right)** The mean activity level for each trial is plotted for DN-A (y-axis) against the mean Difficulty composite scores (x-axis). **(Bottom, Right)** The mean activity level for each trial is plotted for FPN-B (y-axis) against the mean Difficulty composite scores (x-axis) revealing a strong, positive relation. Pearson’s correlation values are shown in the bottom left corners. Scene Construction scores track DN-A activity and Difficulty scores track FPN-B activity, but not vice versa, illustrating a functional double-dissociation between these two closely juxtaposed networks.

Multiple regression confirmed the double dissociation. In a model featuring Scene Construction, Difficulty and Others-Relevant scores, Scene Construction accounted for the most variance in DN-A activity across trials and was the only significant predictor, as described above (R^2^ = 0.36, *p* < 0.001). For FPN-B, Difficulty was the only significant predictor and accounted for most of the model’s variance in network response (R^2^_Difficulty_ = 0.48, *p* < 0.001; R^2^_Full Model_ = 0.50).

To better interpret the robustness of these relations, we also calculated the internal reliability of our measures and estimated the proportion of explainable variance for each composite-network pair. The split-half reliability of the composite scores was high (*r*_Scene Construction_ = 0.92, *r*_Difficulty_ = 0.95), as was reliability of the network activity values (*r*_DN-A_ = 0.87, *r*_FPN-B_ = 0.86^2^). Comparing our models’ R^2^ values to the product of these reliability scores, for each composite-network pair (as an estimate of explainable variance) suggested that Scene Construction scores accounted for about 47% of the explainable reliable variance in DN-A activity, and Difficulty scores for about 60% in FPN-B activity. Overall, the dissociation between DN-A and FPN-B adds to evidence that parallel association networks, with side-by-side regions across association cortex, can be robustly functionally dissociated (as originally suggested by Fedorenko et al. 2012).

### Strategy Composite Score Correlations After Regression of Difficulty

Given evidence of a possible confounding effect of Difficulty on multiple composite-network relations, in *post hoc* analyses, we regressed the Difficulty composite scores from all other composites. Residual data continued to reveal a strong, selective relation between Scene Construction scores and DN-A activity (*r* = 0.59, CI [0.50, 0.68]). Effort does not account for Scene Construction’s relation to DN-A response (see top panel of Figure 12).

**Figure 12.**
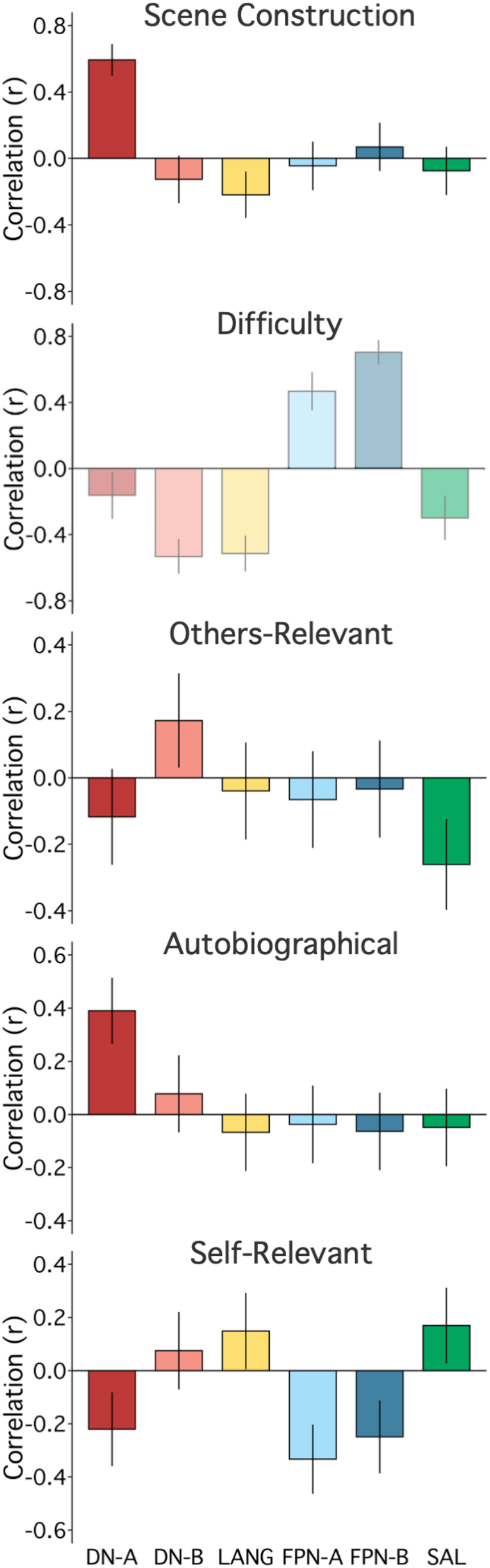
Strategy composites are associated with differential and selective network activity after regression of Difficulty. Given the possibility of a confounding effect of Difficulty on estimates of network selectivity, the network correlation bar plots were recomputed after regressing the Difficulty composite scores (see text). **(Scene Construction)** Scene Construction composite scores maintained a strong and selective correlation to DN-A activity. **(Difficulty)** As mandated by the analysis, variation related to the Difficulty Composite scores was removed. Since correlations could not be estimated to a vector of zeros, the original Difficulty plot (from Figure 8, faded here) is shown for reference. **(Others-Relevant)** The Others-Relevant composite scores maintained a weaker but selective relation to DN-B activity. **(Autobiographical)** Of interest, the Autobiographical composite scores, once Difficulty was regressed, revealed a pattern similar but weaker to that of Scene Construction. **(Self-Relevant)** The Self-Relevant composite scores were non-specific. Networks, from left to right: Default Network A (DN-A, red), Default Network B (DN-B, pink), the Language Network (LANG, yellow), two candidate Frontoparietal Control Networks (FPN-A, light blue; FPN-B, dark blue) and the Salience Network (SAL, green).

Autobiographical scores also showed a selective, positive correlation to DN-A activity after Difficulty regression, and the correlation between Scene Construction and Autobiographical composites strengthened. Scene Construction was still the strongest predictor of DN-A response, including within a model including both Autobiographical and Self-Relevant composites (both of which were also significant: R^2^_Scene Construction_ = 0.25, *p* < 0.001; R^2^_Autobiographical_ = 0.09, *p* < 0.05; R^2^_Self Relevant_ = 0.05, *p* < 0.001; R^2^_Full Model_ = 0.40). We unpacked these results in further *post hoc* testing (see *Further Tests Do Not Support Contribution of DN-A to Recollection of the Personal Past*, below).

Others-Relevant scores maintained a weak but selective positive relation to DN-B response after regression of Difficulty. We initially expected a relation between Others-Relevant scores and DN-B response, based on evidence of DN-B recruitment for social functions (e.g., DiNicola et al. 2020) and ample evidence that relevant regions participate in representing others’ thoughts (e.g., see Lieberman 2007, Koster-Hale & Saxe 2013, Schurz et al. 2020). A model featuring the four remaining composites was significant (*p* < 0.01) but only modestly predicted variance in DN-B activity overall (R^2^_Full Model_ = 0.08), and the positive relation to Others-Relevant scores accounted for similar variance to a negative relation to Scene Construction (R^2^_Others-Relevant_, = 0.03, R^2^_Scene Construction_ = 0.03).

For both Autobiographical and Self-Relevant composites, other interpretable network correlations did not survive regression of the Difficulty composite scores (compare Figure 8 to Figure 12). These results aligned with the evidence of multicollinearity between Difficulty, Autobiographical and Self-Relevant scores in our initial regression model. Comparing Autobiographical and Self-Relevant scores to mean behavioral RT also supported a Difficulty confound: both composite scores were strongly, negatively correlated with RT (*r* = −0.44 for both). No relation to RT was found for Scene Construction composite scores (*r* = 0.00) and a much weaker relation for Others-Relevant scores (*r* = −0.16).

These findings indicate that our data could reduce to three clusters, related to Scene Construction, Others-Relevant, and trial Difficulty dimensions (e.g., with Self-Relevant scores along a continuum of effort). Revisiting our initial composite characterization, both regressing and reverse-coding Difficulty composite strategies (i.e., with higher values for easier and subjective trials) preserved Scene Construction and Others-Relevant correlations, separable from an intercorrelated Self-Relevant cluster.

The relation between Scene Construction scores and DN-A response and the separate relation between Difficulty scores and FPN-B response become even more compelling given these additional analyses. As an additional *post hoc* test, in the next section, we probed these two relations to assess whether maps based solely on trial groupings from the composite scores could yield selective network recruitment within individual participants.

### Distinct Networks Can Be Recapitulated Within Individuals From Small Numbers of Trials That Differ in Strategy Composite Scores

As a test of the discovery that Scene Construction and Difficulty scores differentiate activity between juxtaposed networks, we created within-individual, whole-brain contrast maps using trials with high and low scores on the Scene Construction and Difficulty composites. Importantly, for Scene Construction, we restricted the analysis *only* to trials from the original control conditions. Thus, this contrast is completely orthogonal to the originally envisioned condition contrasts in DiNicola et al. (2020).

Each individual’s estimated DN-A border was overlaid upon the Scene Construction contrast maps. Results revealed alignment between the contrast maps and DN-A (Figure 13, left column). Overlap was observed within midline regions, including in retrosplenial cortex and posterior parahippocampal cortex (PHC), previously linked to Scene Construction ratings (e.g., Andrews-Hanna et al. 2010, see also Hassabis & Maguire 2007), as well as in distributed DN-A regions, including in dorsolateral PFC, lateral posterior parietal cortex and lateral temporal cortex. Although correspondence was not perfect and varied by individual, the overlap with DN-A estimates in a subset of participants was striking, particularly given that this contrast included a relatively small number of control trials. Thus, even when only considering trials explicitly designed to minimize demands on episodic projection, contrasting trials with higher versus lower Scene Construction scores reveals activity across the distributed DN-A network.

**Figure 13.**
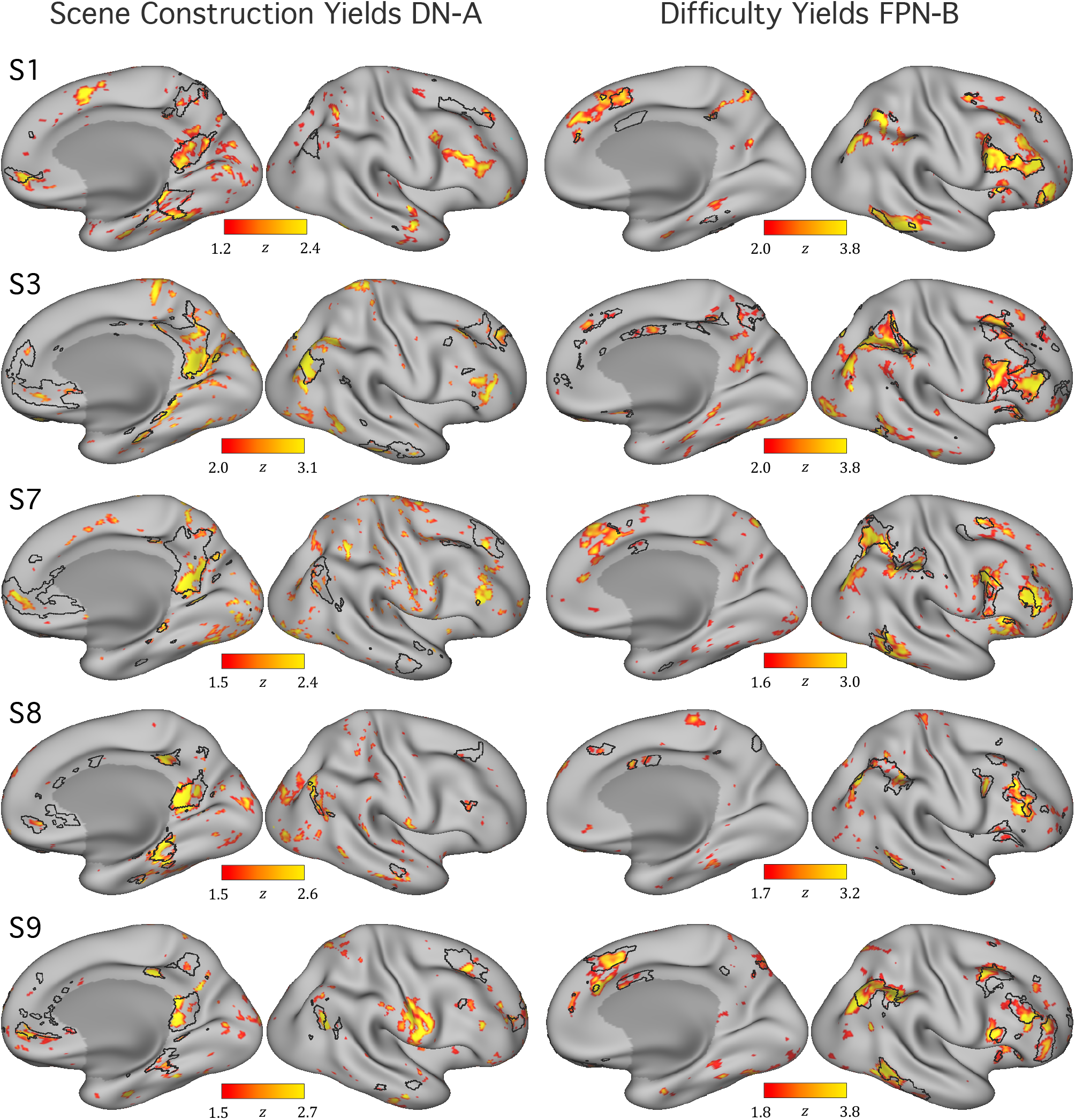
Post-hoc contrasts of extreme trials can recapitulate networks within the individual. As a confirmation of the discovery that Scene Construction and Difficulty are associated with functional MRI response levels in distinct networks, within-individual contrast maps are displayed. **(Left)** For 5 selected individuals, maps illustrate the contrast of the 10 trials with the lowest Scene Construction composite scores versus the 10 trials with the highest scores, selected only from the original control conditions. Note that even with this relatively small amount of data per individual (from trial conditions originally selected to minimize demands on episodic projection) functional MRI differences emerge across the distributed regions that comprise DN-A. The black outlines show the independently estimated boundaries of DN-A for each participant. **(Right)** Maps illustrate the contrast of the 10 trials with the highest Difficulty composite scores versus the 10 trials with the lowest scores. Differences emerge that fall within FPN-B. These contrasts were not possible in initial condition-level analyses (DiNicola et al. 2020) and provide evidence of novel processing insights. Maps are shown for the right hemisphere.

For Difficulty contrasts, each individual’s estimated FPN-B border was overlaid. Comparing the contrast, within individuals, to network borders again revealed evidence of overlap (Figure 13, right column). In most tested individuals,^3^ Difficulty maps corresponded to FPN-B regions across distributed cortical zones, including in prefrontal cortex (as might be expected for cognitive control; e.g., Miller & Cohen 2001; see also Badre & Nee 2018), but also in parietal, temporal, and midline zones. The maps provide evidence that FPN-B supports processes related to effortful control and illustrate the power of the described data-driven approach: individually-defined networks can be reproduced from process-level dissociations, with contrasts created from independent ratings, not possible in prior condition-level analyses. This strategy is powerful even for dimensions (Difficulty) beyond the task’s original design and in networks (FPN-B) not included in the initial analyses.

### Further Tests Do Not Support Contribution of DN-A to Recollection of the Personal Past

In addition to the strong relation between Scene Construction and DN-A, following Difficulty regression, Autobiographical scores showed a selective relation to DN-A response. In additional *post-hoc* tests, we further examined DN-A’s contributions to scene construction and self-reported reliance on mnemonic processes.

Scene Construction and Autobiographical scores were strongly correlated across all trials (*r* = 0.58; Figure 14, top) and even across control trials (*r* = 0.49; Figure 14, middle), presenting a challenge to parsing differences. But among the control trials, subsets showed higher scores on either the Autobiographical or Scene Construction composite (Figure 14, middle). Examining DN-A response across these trials revealed higher values for the Scene Construction subset (*t*(11.40) = 3.24, p < 0.01; Figure 14, bottom).

**Figure 14.**
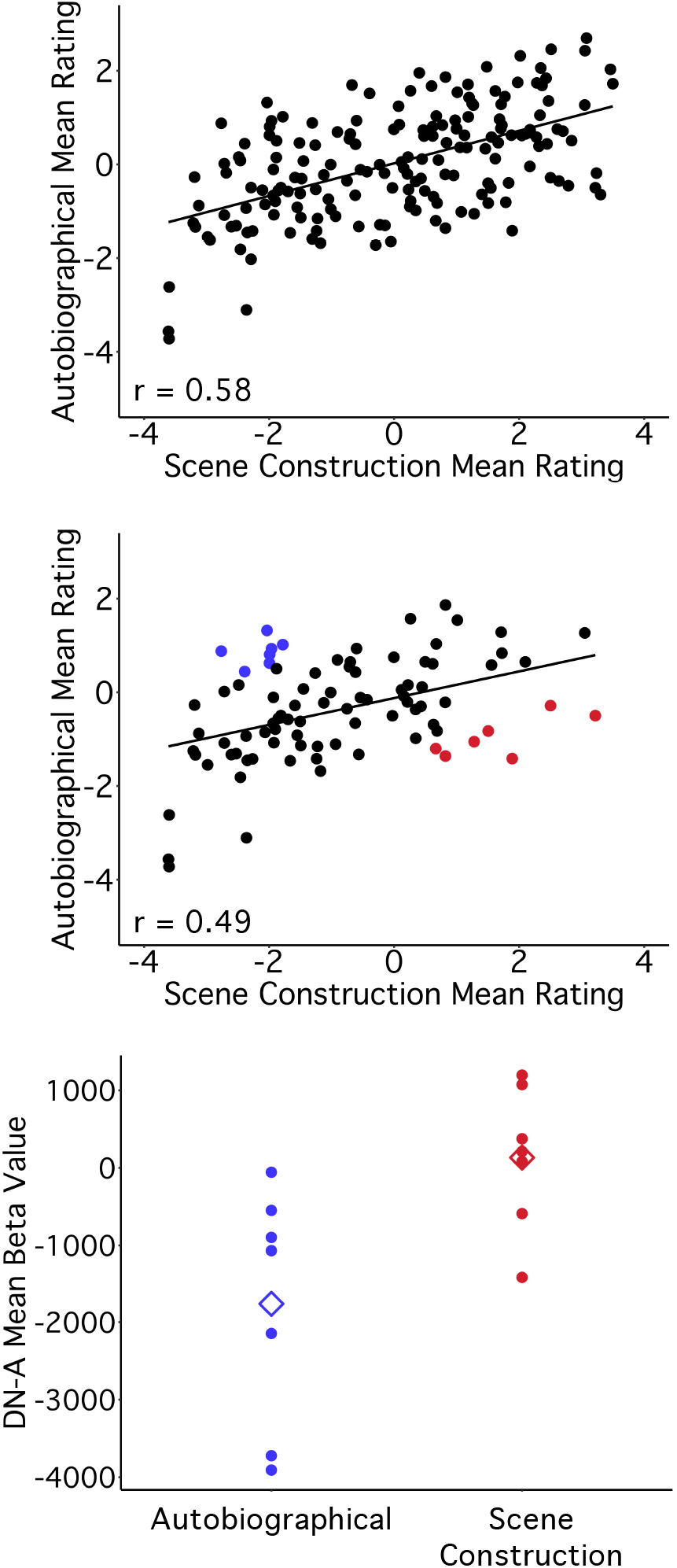
Further exploration of Autobiographical and Scene Construction scores in relation to DN-A response. Following regression of Difficulty, the trial-to-trial correlation between Autobiographical and Scene Construction scores strengthened (*r* = 0.58, **top**). Even for trials originally treated as controls (from Present Self and Past and Future Non-Self conditions), this correlation remained strong (*r* = 0.49, **middle**). But among control trials, subsets of questions were identified that showed higher scores *either* on the Autobiographical (blue) or the Scene Construction composite (red). Plotting the DN-A response values for each trial in these subsets (**bottom**) revealed higher DN-A activity during trials in the Scene Construction subset. Mean DN-A response in each trial subset is shown by a diamond.

What’s more, unpacking the Autobiographical composite, one strategy probe measured reliance on memory (i.e., Pers_Past_Exper in Table 1), while the other measured envisioning a sequence of events (Sequence_Events), also relevant to constructing mental scenes. To assess whether component strategies differentially accounted for variance in DN-A activity, we tested a model featuring each of the Scene Construction and Autobiographical probes in relation to DN-A response (following Difficulty regression).^4^ Within the model (R^2^_Full Model_ = 0.50), three predictors were significant, including both Scene Construction probes (R^2^_Loc_Obj_Places_ = 0.31, *p* < 0.001, R^2^_Visual_Imagery_ = 0.09, *p* < 0.01) and Sequence_Events (R^2^ = 0.10, *p* < 0.01). Little variance in DN-A activity was accounted for by the Pers_Past_Exper probe (R^2^ = 0.01, *p* > 0.05), the only one specific to recollection of the past.

Plotting correlation values between DN-A response and each individual strategy probe (Figure 15, top) illustrated these results, with high correlations to both Scene Construction probes (before and after regression of Difficulty), and higher correlation to Sequence_Events than Pers_Past_Exper within the Autobiographical composite. A positive correlation to Specificity (not assigned to a composite during clustering) was also revealed. Collectively, these results add to evidence for DN-A’s role in constructing mental scenes, likely including specific details of dynamic events.

**Figure 15.**
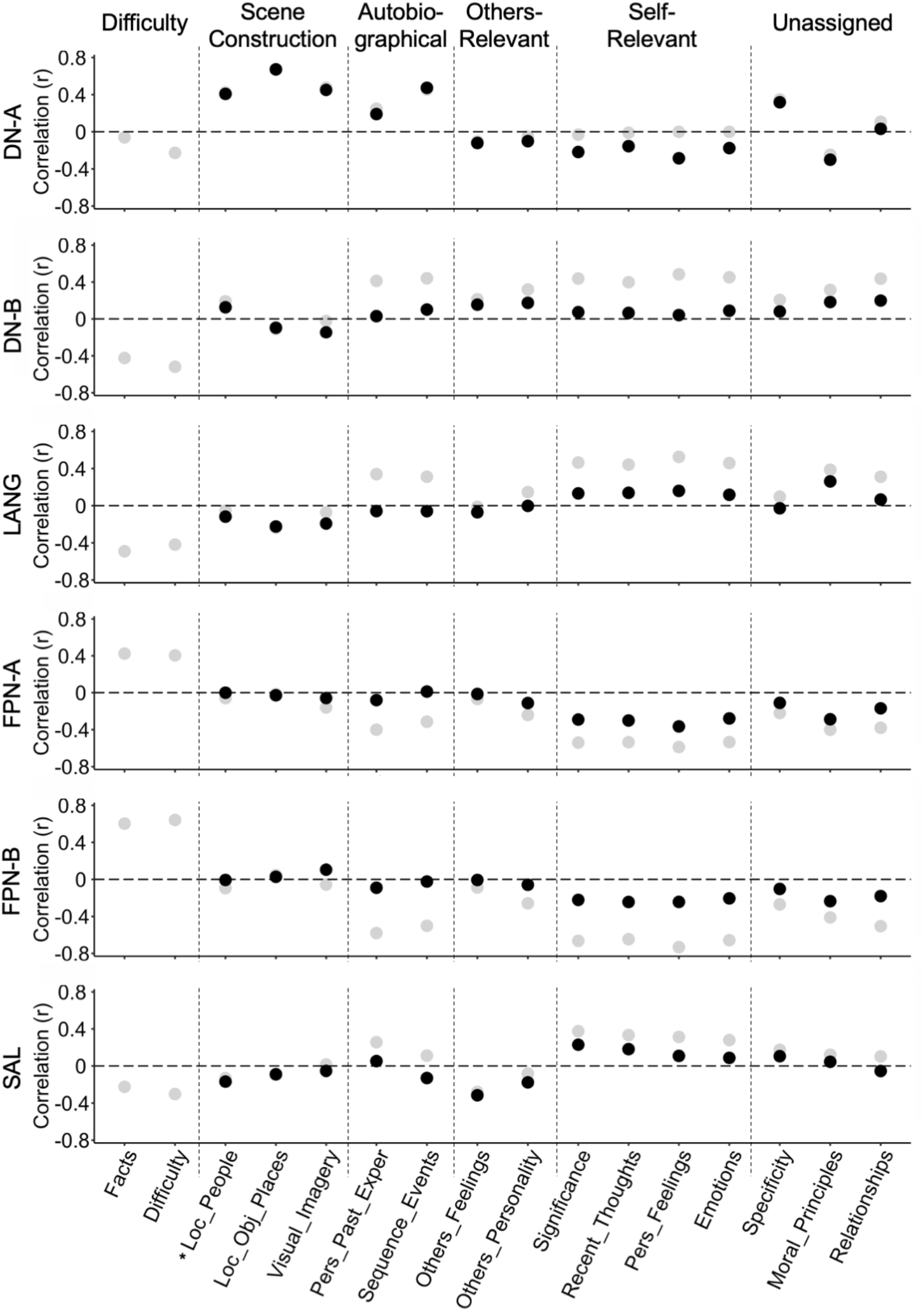
Individual strategy probes for scene construction processes show the strongest relation to DN-A response. For each individual strategy probe from the RSS, trial-level correlations to each network’s response pattern are plotted. Grey circles show correlations prior to Difficulty regression and black circles post-regression. Dotted lines demarcate composites (see Figure 4). The strategy probe for Locations of People (Loc_People, marked with a *) is grouped with the Scene Construction probes for visualization. DN-A (**top**) shows high correlations to Scene Construction probes, as well as to the Sequence_Events probe from the Autobiographical composite. DN-A is more weakly correlated to the Pers_Past_Exper probe. This pattern supports DN-A’s role in mental construction of scenes and events. Patterns across other networks show strong correlations between the FPNs and Difficulty probes (in grey, prior to regression) and a weaker but unique relation between DN-B and Others-Relevant strategies, which survives Difficulty regression. The Difficulty composite comprised Facts and Difficulty strategies, so only preregression correlations to these strategies are shown (in grey).

## Discussion

Processes linked to scene construction selectively recruited one specific distributed network, termed DN-A, that includes PHC, retrosplenial cortex and multiple cortical association regions.^1^ The interwoven but anatomically distinct DN-B showed no such response. The functional dissociation between these two juxtaposed networks was striking (Figure 9) and suggests that DN-A is domain specialized. When functional response properties were examined broadly, across multiple distributed association networks, scene construction was associated only with DN-A response and could be further dissociated from responses tracking cognitive effort. Moreover, the relation of DN-A to scene construction held even for trials designed *not* to include episodic memory demands. DN-A appears to subserve scene construction processes, likely encouraged by - but not limited to - autobiographical memory tasks (see also Hassabis & Maguire 2007, 2009). We discuss the implications of these observations as well as the opportunities and limitations of our methods that leverage trial-to-trial variation in processing demands to constrain understanding of network functions.

### Trial-Level Variation in Scene Construction Robustly Tracks DN-A Response

Comparing trial-to-trial variation in Scene Construction ratings to network activity revealed a selective, strong relation to DN-A response. Prior notions that a monolithic DN makes an extremely broad processing contribution to diverse forms of mental simulation (e.g., Buckner & Carroll 2007, Spreng et al. 2009) are not consistent with our data. Rather, our results support the hypothesis that DN-A and DN-B contribute to distinct domains of processing, and further, that the hippocampally-linked DN-A specifically subserves scene construction, including as used during episodic remembering and imagining future scenarios, but also during atemporal imagination and for scenarios that are not necessarily relevant to oneself (see Hassabis & Maguire 2007, 2009, Andrews-Hanna et al. 2010). The selectivity for processes associated with scene construction and not episodic retrieval is important to refining functional understanding.

DN-A is strongly recruited, on average, for trials featuring remembering and constructing future scenarios when these trials are contrasted to control trials involving semantic or non-personal reference (DiNicola et al. 2020; see also Addis et al. 2007, Spreng et al. 2009). But such complex task trials rely on multiple distinct component processes (for relevant discussion, see Hassabis & Maguire 2009). While mental scene construction presumably always utilizes some form of internal process as the scenes are imagined (not experienced), a clear finding is that DN-A activity is not specifically linked to whether a trial demands reliance on episodic memory - that is, retrieval from one’s own personal past. Within the entire trial set, the relation between DN-A and reported utilization of personal past experiences was weak, and when restricted to control trials, negative. However, trials, including controls, that rated low on utilizing past personal experience activated DN-A to the degree they rated high on Scene Construction. Thus, the high average response of DN-A across trials involving episodic remembering and imagining future scenarios appears driven by covariation with the more basic component process of scene construction. To the degree a trial encouraged participants to construct a mental scene with vivid imagery and awareness about spatial locations of objects or places, the response in DN-A increased. Providing further support that scene construction is the core process driving DN-A activity, contrast maps made *only* from control trials, using ratings on Scene Construction probes, overlapped with DN-A estimates within individuals (Figures 13).

A nuance to interpreting our data arose in the more detailed analysis of behavioral strategies once Difficulty was regressed. Although Autobiographical ratings also tracked DN-A response, a strategy probe for envisioning event sequences largely accounted for this relation. The probe directly measuring reliance on personal past experiences still only weakly related to DN-A activity. Across all probes, those for visualizing scene and event details, even including the specificity of such details, related most strongly to DN-A. This strategy pattern supported DN-A’s role in internally constructing scenes, including dynamically unfolding event sequences, even when minimally reliant on the personal past (see also Hassabis et al. 2009).

The observed relations between DN-A and scene-relevant probes were not only strong but selective. Comparing functional relations across multiple distributed association networks illustrated that *only* DN-A was recruited for Scene Construction. Interwoven DN-B showed almost no relation to scene-relevant strategies, nor did any of four other juxtaposed networks. And although multiple nearby networks tracked Difficulty (a composite measuring cognitive effort), DN-A showed no relation to Difficulty scores, and Difficulty-related networks did not track Scene Construction. DN-A can thus be functionally dissociated from parallel - and even interdigitated - networks within association cortex through a unique role in scene construction.

These results build evidence in support of domain specialization for individually-defined, distributed network DN-A. Our findings converge with work linking medial temporal lobe DN regions to constructing mental scenes (Andrews-Hanna et al. 2010; see also Axelrod & Rees 2017, Palombo et al. 2018) and to vivid visual imagery of events (e.g., Wen et al. 2020; Lee et al. 2021). And our work aligns with prior studies linking regions of DN-A to category-specific reasoning about places, including in posteromedial cortex (Peer et al. 2015, Silson et al. 2019, Woolnough et al. 2020) and in more distributed regions (Deen & Friewald 2020). Ultimately, rather than the canonical DN, our evidence supports DN-A contributing to a previously-hypothesized ‘construction system’ (Hassabis & Maguire 2009), supporting processes for mentally creating coherent scenes.

### Method Intuitions: Behavioral Ratings Capture Stable Properties of Trial-to-Trial Variation

In addition to network insights, the present work provided evidence that behavioral strategy ratings can tap into stable trial-level properties. If behavioral ratings varied by respondent group or by individual answers to trial questions, they would not inform independent neuroimaging data. But across two behavioral experiments, ratings of strategy probes showed striking stability. Even for individual trials, patterns were highly reliable, akin to trial-level fingerprints, and inter-experiment reliability was high for nearly every strategy probe (Figures 1 and 2). What’s more, correlated strategies formed replicable clusters, suggesting that, for trials like ours (requiring consideration of different scenarios), participants have insight into *how* they choose a response (Figures 3 and 4). Self-reported strategy ratings thus capture stable trial properties, informative to independent neuroimaging results from the same trial set (see also Andrews-Hanna et al. 2010).

Here, neuroimaging data were also reliabile,^2^ with activity levels accounting for behavioral variation. Specifically, trial-to-trial behavioral strategy ratings correlated with fMRI response activity within the individually-identified networks. At first glance, our correlation results are surprising. The scatterplot showing Scene Construction scores plotted against DN-A response, for example, reveals a remarkably clean relation (Figure 9), reminiscent of the kind of artifactual relations that emerge when there is circularity in region definition, colloquially known as double dipping (Kriegeskorte et al. 2009; Vul et al. 2009). The network regions analyzed here were defined a priori without any bias to elicit well-behaved relations. The well-behaved plots likely emerge because of the stability of the estimates. Each dot in the plots depicted in Figures 9 and 11 represents not a single *person* but two robust estimates - two means of reliable datasets - for a single *trial*. Each dot, therefore, includes a lot of data, from multiple groups of participants, stabilizing the estimate.

### Limitations and Considerations for Future Work

Despite the strengths of this approach, our datasets were limited by the trials and strategy probes we used. We asked online participants to rate strategy probes for multiple questions in a row, so we restricted the total number of probes. Additional ratings could target such dimensions as perspective, temporal orientation, emotional valence, or certainty (e.g., Andrews-Hanna et al. 2013, Stawarczyk et al. 2013). Our task could also be expanded to include additional trial questions. Trials were not explicitly designed to target social reasoning or effort, for example, which impacted our ability to probe relevant processes.

Varying levels of cognitive effort across trials appears particularly crucial to dissecting network functions. In the present work, a confounding effect of Difficulty impacted the Autobiographical and Self-Relevant composites. As discussed elsewhere, failure to control for difficulty can lead to spurious interpretations of differences between other trial features (e.g., see Caramazza & Shelton 1998). Our task was designed to target other dimensions but nonetheless varied in difficulty, and our findings leave open questions about potential contributions of individually-defined networks to complex affective, narrative, and other processes that could relate to Self-Relevant probes, when difficulty is better controlled.

The described composite-network links also raise questions that could not be answered with the current data. For example, though we largely focused here on the strongest network-composite relation relevant to Difficulty (i.e., FPN-B), *both* frontoparietal networks (FPN-A and FPN-B) showed positive correlations to the Difficulty composite score. A question for future work concerns the precise roles of these networks. Distinct contributions to aspects of control and to network coordination have been proposed (see Dixon et al. 2018, Badre & Nee 2018, Murphy et al. 2020, Nee 2021, see also Marek & Dosenbach 2018) and yet-unappreciated roles are also a possibility (e.g., even beyond the control domain).

In addition, the observed dissociation between DN-A and FPN-B supports a hypothesis that tightly-juxtaposed association networks can distinctly subserve more domain-specialized (DN-A) or domain-general (FPN-B) processes (see also Fedorenko et al. 2012, DiNicola & Buckner 2020). Whether these patterns hold across all distributed association zones is an open question. Contrast maps produced from strategy ratings showed network overlap not only in specific regions, but across distributed cortical zones for both DN-A (in relation to Scene Construction) and FPN-B (to Difficulty; Figure 13). These maps, along with findings of task differentiation in multiple network zones (DiNicola et al. 2020; see also Deen & Friewald 2020), lead us to predict that functional distinctions span the cortex. We aim to test this hypothesis more directly in future work.

Finally, although DN-A has been linked to the hippocampal formation (Braga et al. 2019) and the hippocampus has long been shown to play a role in representing space (e.g., O’Keefe & Dostrovsky 1971) and scene construction (e.g., Hassabis & Maguire 2007; see also Maguire et al. 2016), the present analyses were limited to cortical network regions. Further examination of network connectivity to and function of the hippocampus itself could further clarify DN-A’s functional role.

## Conclusions

Processes linked to scene construction selectively recruited DN-A. Even when examining trials explicitly designed *not* to rely on personal past experiences, neuroimaging contrasts created from extreme Scene Construction ratings revealed a preferential response in DN-A. Scene Construction did not recruit interwoven DN-B or four other networks, and DN-A response did not track Difficulty. These results suggest that parallel distributed networks in association cortex are functionally distinct with DN-A making a domain-specialized processing contribution that can be robustly functionally dissociated from multiple other association networks.

## Acknowledgments

We thank T. Konkle and P. Mair for providing crucial statistical insights. A. Youssoufian, H. Becker, K. Miclau and A. Song assisted in neuroimaging data acquisition and stimuli generation. We thank the Harvard Center for Brain Science neuroimaging core and FAS Division of Research Computing. We thank T. O’Keefe, H. Hoke and R. Mair for assisting in neuroimaging preprocessing optimization. The multi-band EPI sequence was generously provided by the Center for Magnetic Resonance Research (CMRR) at the University of Minnesota. We thank J. Andrews-Hanna for supplemental task and strategy probe stimuli and insights. We thank Hayley Dorfman and Hanna Hillman for insights into online data collection.

## Footnotes

1 The labels DN-A and DN-B were chosen because ofthe literature's description of the broader network as the default network or default mode network. DN-Aand DN-B are not subnetworks of the default network. Rather, by our estimates, they are two parallel networks that have been historically mischaracterized as a composite monolithic network because of spatial blurring in group-averaged data (including in our own work; e.q., Buckner et al. 2008). Thus, the labels DN-A and DN-B are labels of convention to relate our present description of these parallel distributed networks to the historical description of the default network. The network labels should not be taken to imply strong assumptions about the functional domains of these networks.

2 Split-half reliobility for all composite scores was greater than 0.90 and for all networks was greater than 0.70, except for the SAL network. One hypothesis for lower SAL reliability (0.52) in this task is individual differences, across trials, in which elements captured attention, but this requires testing.

3 For one individual (S4), the Difficulty contrast map showed more overlap with an estimate of FPN-A. As described previously, this individual had a more ambiguous k-means output (S11 in DiNicola et al. 2020), and this result likely reflects uncertainty in network assignment, rather than true network differences, in this subject. This result raises the importance of exploring network responses in additional individuals.

4 Within this model, the two Scene Construction probes were highly intercorrelated (VIF > 3). A similar model, with these probes combined, produced comparable results. Most DN-Avariance was captured by Scene Construction (R^2^_Scene Construction_ = 0.25, P < 0.001; R^2^_Full Model_ = 0.40) followed by Sequence_Events (R^2^ = 0.12, P < 0.001), and almost none by Pers_Past_Exper (R^2^ = 0.02, P <0.05).

